# Voltage imaging identifies spinal circuits that modulate locomotor adaptation in zebrafish

**DOI:** 10.1101/2021.08.20.457089

**Authors:** Urs L. Böhm, Yukiko Kimura, Takashi Kawashima, Misha B. Ahrens, Shin-ichi Higashijima, Florian Engert, Adam E. Cohen

## Abstract

Motor systems must continuously adapt their output to maintain a desired trajectory. While the spinal circuits underlying rhythmic locomotion are well described, little is known about how the network modulates its output strength. A major challenge has been the difficulty of recording from spinal neurons during behavior. Here, we use voltage imaging to map the membrane potential of glutamatergic neurons throughout the spinal cord of the larval zebrafish during fictive swimming in a virtual environment. We mapped the spiking, subthreshold dynamics, relative timing, and sub-cellular electrical propagation across large populations of simultaneously recorded cells. We validated the approach by confirming properties of known sub-types, and we characterized a yet undescribed sub-population of tonic-spiking ventral V3 neurons whose spike rate correlated with swimming strength and bout length. Optogenetic activation of V3 neurons led to stronger swimming and longer bouts but did not affect tail-beat frequency. Genetic ablation of V3 neurons led to reduced locomotor adaptation. The power of voltage imaging allowed us to identify V3 neurons as a critical driver of locomotor adaptation in zebrafish.

## Introduction

Dynamically adapting motor output is an essential function of vertebrate locomotor control. Animals must adjust the force of their muscles to change speed or to respond to changes in internal or external forces. The core networks that generate the rhythmic activity essential for locomotion have been extensively studied in many model systems (Grillner and Manira, 2015). Much less is understood about how these networks are dynamically modulated. One technical challenge is the difficulty of spinal cord recordings during behavior. Rodent spinal cords are hard to access optically or electrically *in vivo*, and organisms more amenable to optical methods such as *Xenopus* tadpoles or zebrafish larvae have dynamics faster than the temporal resolution of Ca^2+^ imaging. Thus population-wide measurements of membrane potential in locomotor circuits have so far only been possible in invertebrate preparations using voltage-sensitive dyes (Bruno et al., 2015; Tomina and Wagenaar, 2017).

Voltage imaging using genetically encoded voltage indicators (GEVIs) has the potential to overcome these limitations. Recent advances in sensors (Abdelfattah et al., 2019; Adam et al., 2019; Chien et al., 2021; Gong et al., 2015; Piatkevich et al., 2019; Villette et al., 2019), optics (Fan et al., 2020), and analysis tools (Cai et al., 2021; Xie et al., 2021) have made it possible to record subthreshold and spiking activity from multiple cells *in vivo*. These advances open the possibility to map voltage dynamics across a population and to characterize their activity in an unbiased way.

At five days post fertilization (dpf), zebrafish larvae swim in discrete bouts and show a range of behaviors, including fast and slow forward swimming, turns, escape responses and prey capture (Budick and O’Malley, 2000; Marques et al., 2018). Changes in locomotor speed are associated with changes in tail beat frequency in the range from 20 to 70 Hz. Distinct populations of premotor excitatory interneurons are recruited at different tail beat frequencies (Ljunggren et al., 2014; McLean et al., 2007, 2008; Wahlstrom-Helgren et al., 2019). However, adapting motor output can also happen via changes in tail amplitude or force, without substantial changes in frequency (Ahrens et al., 2012; Kawashima et al., 2016; Portugues and Engert, 2011), and a larval zebrafish can modulate its average velocity by changing the fraction of time it is swimming vs. resting. The spinal mechanisms underlying adaptation in swim force and bout duration are currently unknown.

Here we use light-sheet voltage imaging to map neural activity in the zebrafish spinal cord during slow forward swimming in a virtual environment. We characterized in detail the patterns of network activity and sub-cellular action potential propagation across all glutamatergic sub-types. In a previously uncharacterized cell type, spinal V3 interneurons, we found that activity was correlated with swimming but in a non-rhythmic way. V3 neurons are defined by sim1 expression and in mice V3 neurons make connections to motor neurons both ipsi- and contralaterally (Chopek et al., 2018; Zhang et al., 2008). Loss of function and optogenetic experiments in mice suggested a role of V3 neurons in left-right coordination of locomotory activity (Danner et al., 2019; Zhang et al., 2008).

We found that V3 neuron activity during swimming correlated with the strength and duration of each swim bout. We used optogenetic activation and genetically targeted ablations of V3 neurons to characterize the functional role of V3 neurons in larval zebrafish and showed that V3 neurons act as a modulator of swimming, primarily by affecting swim strength and bout length. These findings revealed a new mechanism by which spinal networks modify locomotion and demonstrate the power of voltage imaging to uncover previously inaccessible neural dynamics.

## Results

### Distinct oscillatory and non-oscillatory sub-populations of excitatory spinal neurons

To evoke patterns of neural activity associated with naturalistic swimming in immobile fish, we implemented virtual reality feedback (Ahrens et al., 2013) in a custom-built light sheet microscope (Fig. 1A, S1). Fish were paralyzed with 1 mg/ml α-bungarotoxin and placed in the recording chamber. Electrical recordings from a ventral nerve root (VNR) on one side of the fish were processed in real time to calculate a fictive swim strength, which depended on the VNR amplitude and frequency (Methods). In order to generate robust swim responses we created a virtual reality (VR) environment, and we emulated a condition where the animal was swept backwards by a constant water current. Specifically, the swim strength was scaled by a user-selected feedback gain to obtain a fictive swim speed, and then subtracted from the motion of a drifting grating projected from below. This setup provided independent control of the baseline grating drift speed and the feedback gain and helped to invoke consistent fictive swim bouts interspersed with rests. Fictive swimming produced average cycle frequencies per bout of 23 ± 4 Hz (mean ± s.d., *n* = 840 bouts, 9 fish), an average bout duration of 0.4 ± 0.4 s and average inter-bout intervals of 0.8 ± 0.8 s.

**Figure 1.**
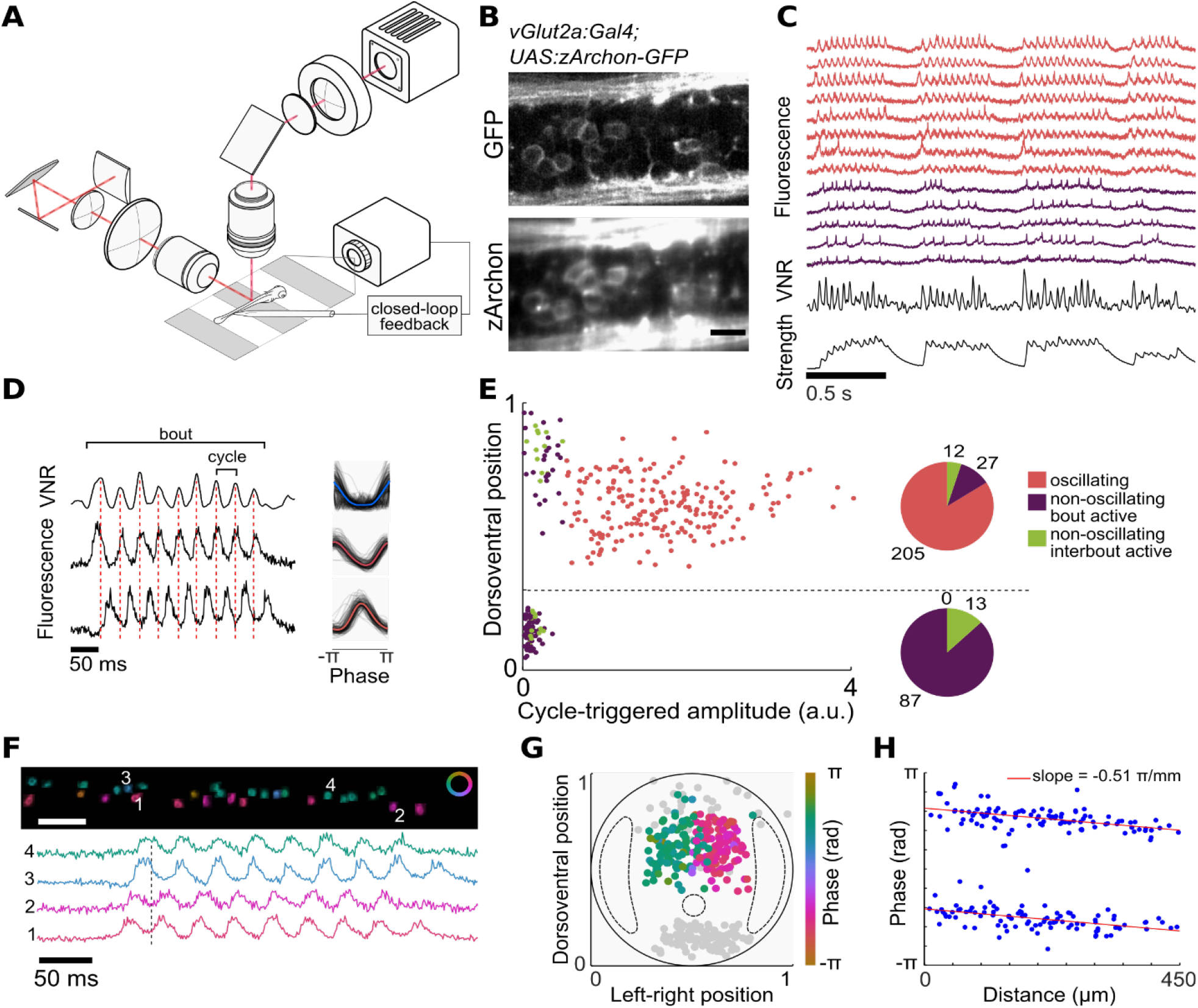
Voltage imaging in spinal neurons during fictive swimming. (A) Schematic of the light-sheet microscope, ventral nerve root recording, and closed loop visual feedback. (B) zArchon1-GFP expression in the *Tg(vGlut2a:Gal4; UAS:zArchon1-GFP)* transgenic line. Top: Two-photon image of GFP expression marker; Bottom: light-sheet image of zArchon1 fluorescence in the same focal plane. Scale bar 10 μm. (C) Fluorescence traces showing *n* = 13 simultaneously recorded neurons, divided into oscillating (red) and non-oscillating (purple) activity patterns. Bottom: processed VNR signal and derived swim strength. (D) Left: VNR signal and fluorescence of two simultaneously recorded oscillating cells. Right: VNR cycle-triggered averages. (E) Relationship between cycle-triggered amplitude and dorso-ventral position for *n* = 344 neurons, 9 fish. The pie charts show the distribution of activity types in the dorsal and ventral subpopulations. (F) Phase map of simultaneously recorded oscillating neurons. Cell bodies are colored according to their phase relative to the VNR signal. Scale bar 50 μm. (G) Transverse view showing cell body positions of oscillating neurons, color coded by phase relative to VNR. Non-oscillating cells in gray, dotted lines indicate position of the lateral neuropil region and the central canal. (H) Relationship of average phase and cell body position along the rostro-caudal axis. The two populations indicate cell bodies on the left and the right side of the spinal cord. The slope indicates the phase delay along the tail. Slope = -0.51 ± 0.06 π radians/mm, r^2^ = 0.96.

To visualize neural dynamics in fish performing fictive swimming, we generated the transgenic line *Tg(UAS:zArchon1-GFP)* to express the voltage sensor zArchon1 (Piatkevich et al., 2018). We then used the *Tg(vGlut2a:Gal4)* driver line (Satou et al., 2013) to drive expression of zArchon1 in glutamatergic spinal neurons of 5-6 dpf zebrafish. Sensor expression in the transgenic line showed excellent membrane localization and closely resembled previous reports with transient expression (Piatkevich et al., 2018) (Fig. 1B). We used wide-field light-sheet imaging to record the membrane voltage of the glutamatergic neurons at a 1 kHz frame rate, while larvae performed fictive forward swims. We recorded routinely from >10 neurons in parallel (11 ± 7 active cells per field of view, mean ± s.d., max: 29, N = 339 cells, 9 fish) (Fig. 1C).

The voltage waveforms broadly fell into two classes: in some neurons, membrane potential oscillated at constant phase relative to the VNR signal during swim bouts (Supplementary Movie 1); in others, membrane potential showed clear spiking but did not oscillate (Fig. 1C, Supplementary Movie 2). To categorize the neurons and quantify the amplitude and phase of the subthreshold oscillations, we used the VNR signal as a time-base and calculated the VNR-triggered average membrane voltage during a swim cycle (Fig. 1D). Neurons with oscillatory subthreshold voltages were clustered in the middle of the spinal cord along the dorso-ventral axis, above the central canal (Fig. 1E). The phases of the oscillating neurons fell into two groups 180° phase-shifted relative to each other and localized to the left and right halves of the spinal cord, respectively (Fig. 1F, G).

Simultaneous voltage imaging of multiple neurons along a 480-μm long segment of the spinal cord revealed a position-dependent phase delay with a mean slope of 0.51π radians/mm (or 0.087 m/s at an average cycle frequency of 22.6 Hz), corresponding to a wavelength of 3.9 mm (Fig. 1H). At 5-6 dpf, we measured a muscle segment size of 93 ± 8 μm (mean ± s.d.), implying a phase delay of 2.4 ± 0.2% per segment. Previously published VNR recordings reported a similar phase delay of motor output (2.6 ± 1.7% per segment, Masino and Fetcho, 2005). The location and dynamics of the oscillatory neurons indicate that these neurons are primarily V0v (MCoD) and V2a (CiD) premotor excitatory interneurons, which have been reported to show oscillatory dynamics similar to our observations (McLean et al., 2008). These neurons have a spike width of approximately 1 ms (McLean et al., 2008), below the Nyquist frequency of our recordings. Consequently, spikes were only detectable in the most strongly expressing cells (Fig. S2).

The second population consisted of distinctively non-oscillating neurons. This population comprised a subpopulation clustered at the very ventral margin of the spinal cord, and a more scattered subpopulation at the dorsal margin (Fig. 1E, S3). We subdivided the non-oscillating neurons based on whether their spike rate was higher during swim bouts or between swim bouts (Methods, Fig. S3). Most of the ventral population spiked more during swim bouts while the dorsal population contained relatively more neurons that spiked more during interbout intervals (Fig. 1E). Due to their clear anatomical localization and distinctive firing patterns, we focused further studies on the ventral sub-population. Based on their location in the spinal cord and glutamatergic phenotype, we identified these as V3 (VeMe) neurons which have previously been morphologically described as very ventral glutamatergic neurons with a descending axon (Hale et al., 2001; Higashijima et al., 2004) and suggested to be homologs of mammalian V3 interneurons (Goulding, 2009).

### V3 neuron spiking, subthreshold activity, and morphology

We used our extensive dataset of membrane voltage recordings to further characterize the behavior of the V3 neurons. At the start of a swim bout, V3 neurons rapidly depolarized and the spike rate ramped up during the first swim cycle (time constant for spiking onset: τ_on_ = 65 ± 4 ms, Fig. 2A). At the end of a swim bout the VNR signal ended abruptly, but the mean V3 firing rate gradually decreased, starting before the end of the swim bout, without any apparent discontinuity at the end of swimming (τ_off_ = 137 ± 18 ms). The subthreshold voltages also showed fast depolarization at the start of a swim bout and gradual repolarization at the end (Fig. 2A). In contrast to the oscillating dorsal population, neither the subthreshold voltage (Fig. 1E) nor the spike rate (Fig. 2B) oscillated in synchrony with the VNR during swims.

**Figure 2.**
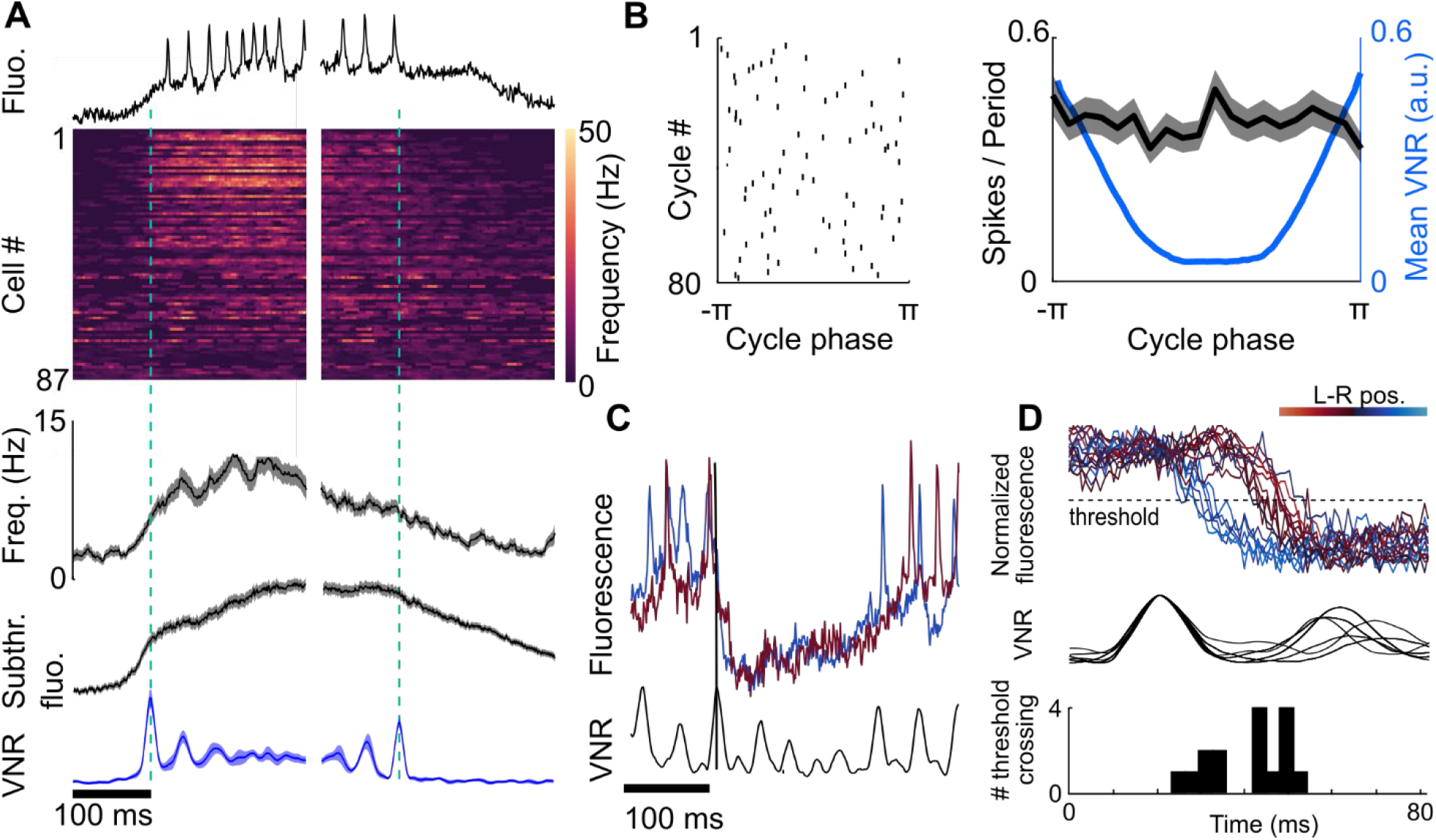
Voltage imaging characterization of V3 neurons. (A) Top: Example V3 recording and spike rates of all V3 neurons aligned to start and end of swim bouts. Spike rate for each neuron was averaged over 5 to 25 bouts. Neurons sorted by bout activity index *I* (Methods). Middle: grand average spike rates for all neurons and subthreshold fluorescence with spike computationally removed. Bottom: Corresponding mean VNR signals aligned to start and end of swim bouts. Shaded area denotes s.e.m. (*n* = 189 bouts). (B) Left: spike raster plot of a single V3 neuron for 80 swim cycles. Spikes were uniformly distributed throughout VNR phase. Right: VNR-triggered grand average spike rate of V3 neurons showing no phase-dependent modulation in spike rate. (C) Rapid inhibition of two V3 neurons on the left (red) and right (blue) sides of the spinal cord during a swim bout. (D) Top: Mean fluorescence transient during mid-bout inhibitory events, aligned relative to preceding VNR peak and color coded according to the cell body position on the left-right axis (*n* = 16 cells, 5-18 events per cell). Middle: Corresponding mean VNR signal. Bottom: histogram of threshold crossings showing two-peaked distribution separated by half a VNR cycle.

We analyzed the subthreshold dynamics of the V3 neurons to determine whether they were driven by a single source or if they comprised distinct sub-ensembles. After digitally removing spikes, we found that the remaining subthreshold dynamics were strongly correlated across cells (Fig. S4). A principal components analysis of *n* = 22 simultaneously recorded V3 cells found that 86% of the variance in the dataset could be accounted for by the first principal component (Fig. S4). These results suggest that the V3 population was primarily driven by a one-dimensional input.

A closer inspection of individual traces, however, revealed some distinctive dynamics in subsets of cells. V3 recordings showed occasional fast membrane hyperpolarizations during swim bouts, indicative of inhibitory inputs to V3 (Fig. 2C). Although V3 spiking and most subthreshold activity were not phase locked to motor rhythm, we noticed that for each cell, the hyperpolarizations bore a definite phase relation to the VNR signal which was constant across multiple events. Moreover, when comparing between cells, the phases of hyperpolarizing events separated into two clusters spaced by half a VNR cycle. These two clusters corresponded to V3 cell body position on the left and right side of the spinal cord (Fig. 2D). Together, these observations imply that V3 neurons receive inputs from an inhibitory population that is phase-locked to the VNR, is sporadically active for one or two swim cycles, and synapses in a lateralized pattern.

High-speed recordings enabled mapping action potential propagation within individual V3 neurons. We applied the sub-Nyquist action potential timing (SNAPT) technique which had previously been used to map action potential propagation in cultured neurons (Hochbaum et al., 2014) (Methods). For each neuron, we extracted spike times from the fluorescence waveform at the soma. We calculated a spike-triggered average movie, typically averaging 100 to 300 spikes from a 25 s recording, and then fit a smoothing spline to the fluorescent trace along the axon and dendrites. This interpolated time trace allowed us to determine the average time-lag of the spike peak at sub-framerate precision (Fig. 3A). In recordings with high signal-to-noise ratio and neurites aligned with the focal plane, this analysis clearly showed action potential initiation at or near the soma and propagation outward (Supplementary Movie 3). The average conduction velocity was 0.19 ± 0.07 m/s (mean ± s.e.m., n = 7 neurons, Fig. 3B, C), roughly two times the speed of the traveling wave in the more dorsal oscillatory population (Fig. 1H) and comparable to the conduction velocity measured for unmyelinated cultured rat hippocampal neurons *in vitro* (Hochbaum *et al*., 2014). To our knowledge, this result provides the first optical *in vivo* measurement of action potential propagation.

**Figure 3.**
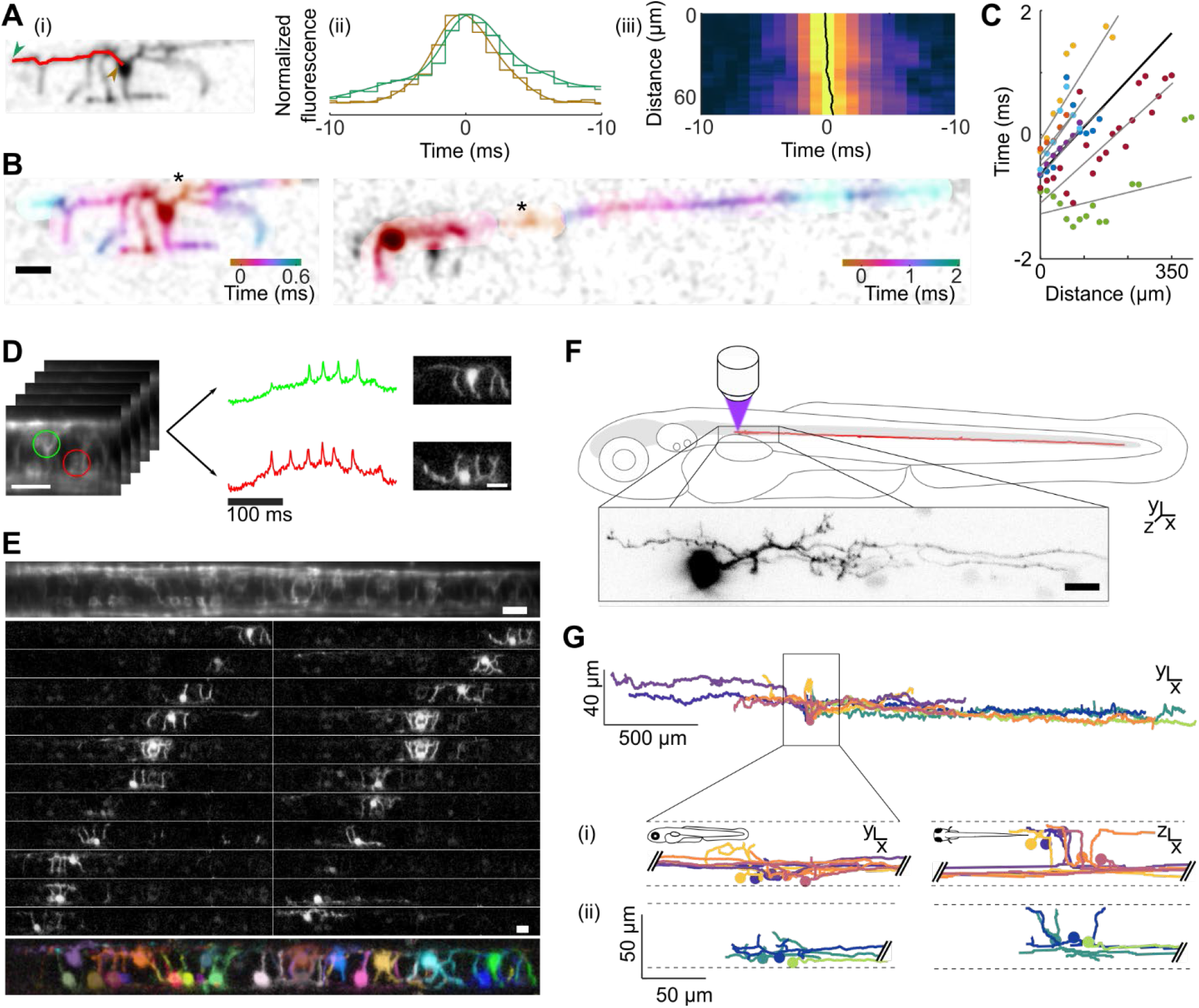
Action potential propagation and V3 morphology. (A) (i) Spatial footprint of a V3 neuron with one neurite marked in red. (ii) Spike-triggered average action potential waveform sampled at discrete 1 ms camera frames (staircase) and interpolated smooth waveforms, from the proximal (brown) and distal (green) ends of the neurite indicated in (i). (iii) Action potential propagation along the neurite marked in (i). Color indicates the normalized fluorescence. Black line indicates the sub-Nyquist interpolated peak timing. (B) Action potential propagation along two neurons. Color denotes time delay, scale bar 20 μm. Action potential initiation sites (black stars) are consistent with action potential initiation at the axon initial segment. (C) Linear fits to the action potential propagation of seven neurons (gray lines) and the average fit (black line). Same colored dots are from the same neuron, each dot represents the binned value over 18 μm or neurite length. The mean conduction velocity was 0.19 ± 0.07 m/s (mean ± s.e.m.). (D) Schematic of activity-based segmentation. Voltage traces measured at the soma were used to calculate an average spike triggered movie and spike waveform. Pixelwise correlation of the spike waveform and the movie revealed the neuron morphology (Methods). (E) Top: Average zArchon1 fluorescence from a voltage imaging movie (25,000 frames). Middle: map of spike-triggered average fluorescence amplitude for each neuron in the recording. Bottom: Composite image with a different color for each functionally identified cell. Scale bar 20 μm. (F) Tracing of a photoconverted V3 neuron in the *Tg(vGlut2a:Gal4; UAS:Kaede)* line. Red line shows extent of axon. Inset shows morphology near the soma. Scale bar 10 μm. (G) Top: morphology of photoconverted V3 neurons. Cell bodies were aligned relative to each other along the rostral-caudal axis and relative to the ventral margin of the spinal cord along the dorso-ventral axis. Bottom: magnified region. Positions were shifted along rostral-caudal axis for better visualization. (i) V3 neurons with bifurcating axons. (ii) V3 neurons with descending axons.

We then used the fact that spike times of V3 neurons were largely uncorrelated between nearby cells to disentangle the signals from densely overlapping neurites. The spike-triggered average movies of each cell clearly showed electrical activation in the cell body and nearby axodendritic arbor, with little contamination from other cells. We calculated a pixelwise cross-correlation of the average spike waveform from each cell with the cell’s spike-triggered average movie. This map revealed the morphology of the neurites around the cell body (Fig. 3D). We computationally extracted the morphology of every neuron, and then re-created a composite image with the individual neurons identified (Fig. 3E).

To explore the morphology of V3 neurons beyond the range accessible via functional unmixing, we tagged the arbors of individual cells using a photoconvertible fluorescent protein. We expressed the protein Kaede in glutamatergic neurons by driving expression of *Tg(UAS:Kaede)* in the *Tg(vGlut2a:Gal4)* line. Single cells were illuminated with a focused laser beam (405 nm) and then photoconverted product was allowed to diffuse through the arbor for 24 h. The morphology from the dendrites around the cell body closely resembled that measured from the functional imaging data (Fig. 3F), validating the activity-based morphological reconstructions.

We observed two morphological classes of V3 neurons, one with a long descending axon and a second one with a bifurcating axon (Fig. 3A). Axons spanned an average distance of 1.7 ± 0.5 mm (mean ± s.d., n = 8 neurons) with bifurcating axons equally split between descending and ascending part (ascending: 0.6 ± 0.4 mm, descending: 0.5 ± 0.2 mm). The population with a descending axon has previously been described as VeMe cells (Hale et al., 2001). The bifurcating neurons represent a previously undescribed morphology. Throughout their extent, the axons of V3 neurons stayed at the ventral margin of the spinal cord (Fig. 3B). A potential target at this position are secondary motor neurons involved in slow swimming (Menelaou and McLean, 2012), suggesting a possible role for V3 neurons in modulating slow swimming.

### V3 neuron activity modulates larval swim bouts during slow swimming

Due to their non-rhythmic spiking, V3 neurons are unlikely to be part of the central pattern generator (CPG) generating the swimming oscillations, yet their distinct firing during swim bouts suggested an acute relationship to locomotion. We hypothesized that V3 neurons might modulate locomotor output.

To test this hypothesis, we recorded V3 spiking activity while changing the feedback gain and the speed of the optic flow in our virtual reality setup to induce swim events of different strengths. This modulation caused V3 neurons to spike during some bouts but not others (Fig. 4A). We measured average fictive swim strength (Methods) as a function of V3 activity. Fish showed a 38% increase in swim strength during bouts in which at least half of the V3 neurons were active compared to bouts where less than half of the V3 neurons were active (swim strength 1.07 ± 0.04 a.u. vs. 0.81 ± 0.05 a.u., mean ± s.e.m., p = 0.0041, paired t-test, n = 9 fields of view from 7 fish, Fig. 4B). We also measured a 50% increase in bout duration in bouts with high vs. low V3 activity (430 ± 48 ms vs. 286 ± 31 ms, p = 0.0135). Remarkably, there was no difference in oscillation frequency between bouts where V3 neurons showed high vs. low activity (22.6 ± 0.8 Hz vs. 22.9 ± 0.9 Hz, mean ± s.e.m., p = 0.4, paired t-test) (Fig. 4B). This observation is in contrast to the activity of many other spinal interneurons whose firing correlates with tail-beat frequency (Ampatzis et al., 2014; Kimura and Higashijima, 2019; McLean et al., 2007, 2008), suggesting a distinct role for V3 neurons.

**Figure 4.**
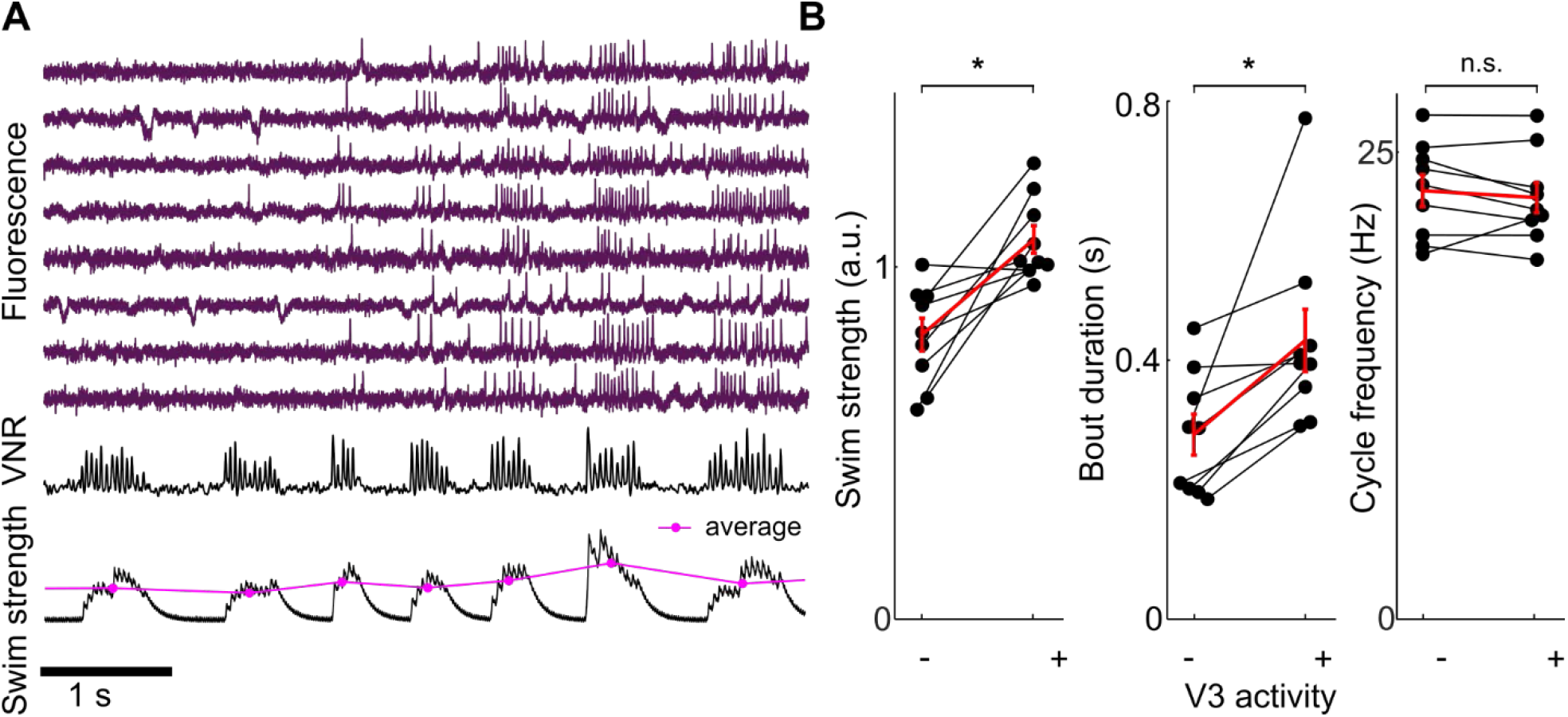
V3 activity correlates with increased swim strength. (A) Eight simultaneously recorded V3 neurons during swims of different strengths. All neurons increased firing during stronger swim bouts. (B) V3 activity correlated with swim strength and bout duration but not cycle frequency. Each dot represents the average of all bouts in each category from one FOV. V3 activity was defined as >50% of V3 neurons in the FOV spiking at least once during the bout. *N* = 9 FOV from 7 fish. Significance threshold α = 0.017 after Bonferroni correction, paired t-test.

We next sought to determine whether the correlation between V3 neuron spiking and bout strength was causal, by optogenetically activating V3 neurons and measuring the effect on tail motions. To that end, we generated a transgenic line conditionally expressing Channelrhodopsin (ChR) WideReceiver (Wang et al., 2009) in glutamatergic neurons (*Tg(vglut2a:loxP-DsRed-loxP-ChR)*) and a second line expressing Cre-recombinase in the ventral p3 domain (*Tg(nkx2*.*2b-Cre)*). This allowed us to express ChR only in V3 neurons (Fig. 5A). To prevent possible visual artifacts during light stimulation and to uncouple the spinal cord circuits from upstream brain areas, we spinalized 5-6 dpf larvae and induced rhythmic tail beating by bath application of NMDA (McDearmid and Drapeau, 2006) (Fig. 5B, C).

**Figure 5.**
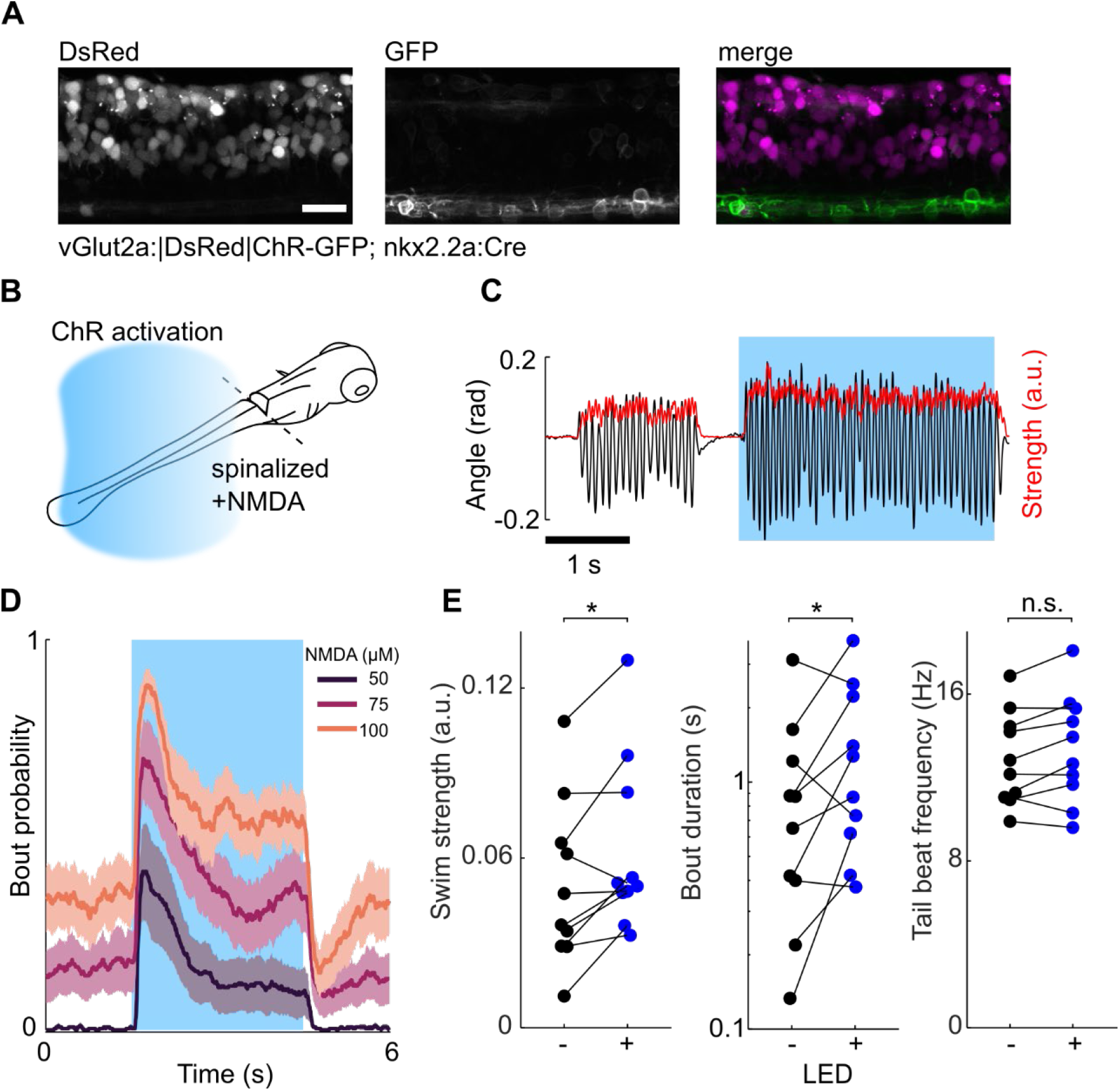
Optogenetic activation of V3 neurons increases swim strength in spinalized larvae. (A) Lateral view of a spinal cord segment of *Tg(vGlut2a:*|*DsRed*|*ChR-GFP; nkx2*.*2:Cre)* larva expressing ChR-GFP in V3 neurons. (B) Schematic of the experimental setup. (C) Spontaneous and optogenetically evoked tail oscillations in the presence of 100 μM NMDA. Blue indicates ChR stimulation. (D) Average bout probability at increasing concentrations of NMDA during ChR stimulation (blue). N = 10 fish, averaged over 20 repetitions. (E) Optogenetic stimulation increased bout duration and swim strength, but not beat frequency. Experiments performed at 100 μM NMDA.

Tonic optogenetic activation of V3 neurons led to an increase in the probability of starting a swim bout, in the duration of the swim bouts, and in the amplitude of the tail oscillations during NMDA induced swim bouts (Fig. 5D). Swim bouts often began immediately after the start of illumination, indicating that V3 activity was sufficient to initiate swimming. Swim bouts also often stopped immediately after the blue light was turned off. At 100 μM NMDA, optogenetic stimulation increased the bout duration, *τ*, (*τ*_*blue*_/*τ*_*dark*_ = 2.0 ± 0.4, mean ± s.e.m., see Methods for regression model and statistical testing) and also increased the swim strength, *S*, (*S*_*blue*_/*S*_*dark*_ = 1.5 ± 0.2, Fig. 5E). In concordance with our data from fictive swimming, we did not observe a significant optogenetically induced change in tail beat frequency, *F*, (*F*_*blue*_/*F*_*dark*_ = 1.04 ± 0.02).

Finally, we tested the effect of V3 activity on freely behaving larvae. We generated a conditional knockout line that expressed the cytotoxic peptide diphtheria toxin A (DTA) under the control of the *sim1a* promoter (*Tg(sim1a:loxP-DsRed-loxP-DTA)*), a marker of V3 neurons (Borowska et al., 2013). To selectively ablate V3 neurons in the spinal cord only, we crossed *Tg(sim1a:loxP-DsRed-loxP-DTA)* to the spinal cord specific Cre driver line *Tg(hox4a/9a:Cre)* (Kimura and Higashijima, 2019). This combination resulted in the loss of >80% of V3 neurons in the spinal cord in 5-6 dpf larvae (Fig. 6A). To induce different swimming speeds, we placed larvae in a 76.2 × 15.7 mm arena and elicited an optomotor response (OMR) by presenting a moving grating at different speeds while monitoring behavior with a high-speed camera (Fig. 6B).

**Figure 6.**
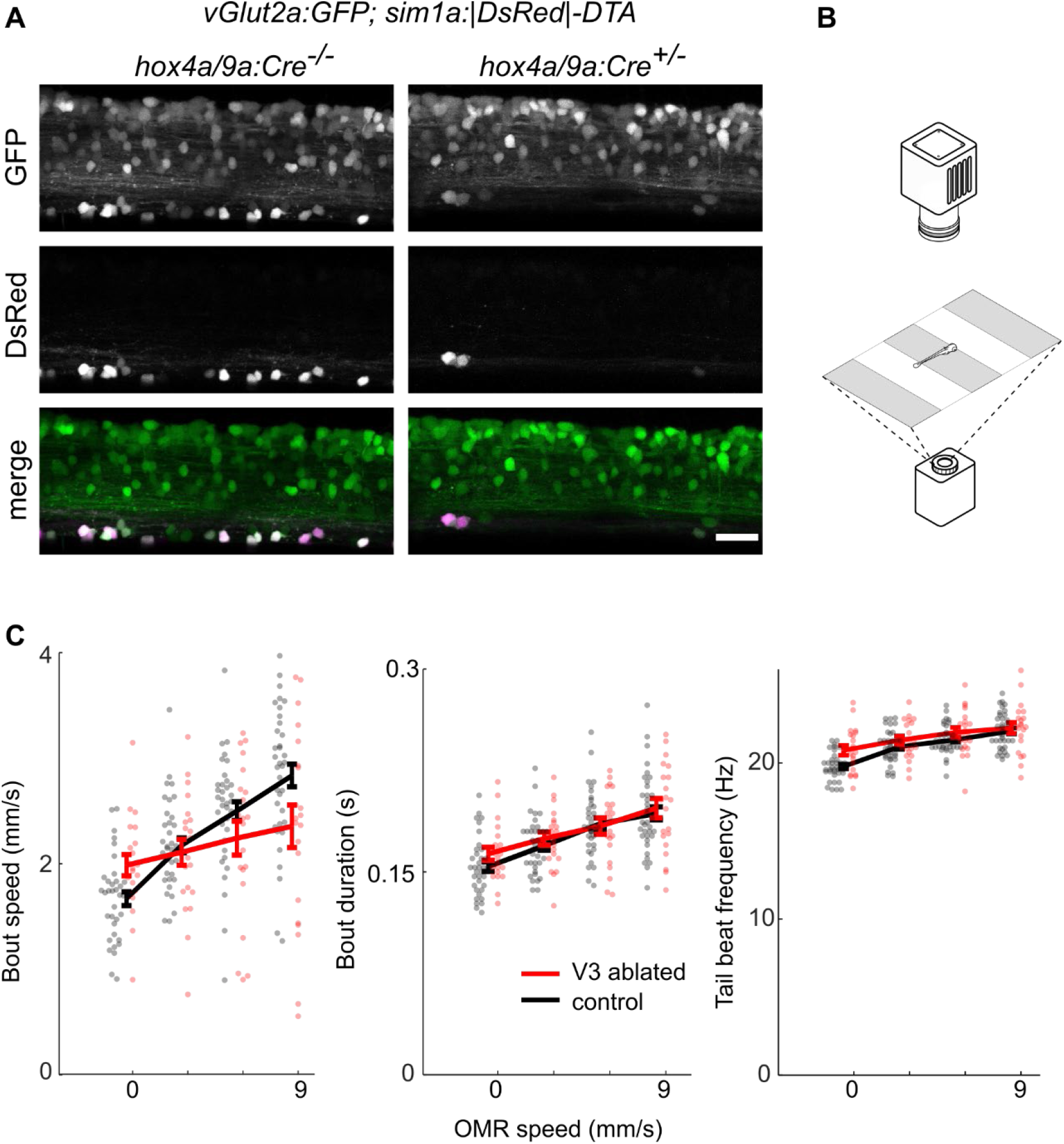
V3 neurons are necessary for swim speed adaptation. (A) lateral view of a spinal cord of *Tg(vGlut2aGFP; sim1a:*|*DsRed*|*-DTA)* without (left panel) or with (right panel) *Tg(hox4a/9a:Cre)*. Cre expression led to the loss of most V3 neurons as seen from the absence of DsRed-expressing cells in the ventral spinal cord. (B) Schematic of the behavioral testing setup. Free-swimming larvae were presented with a moving grating from below and recorded from above. (C) V3 ablated larvae showed reduced modulation of bout speed in response to changes in OMR grating speed, but no difference in tail beat frequency or bout duration. Each dot represents the median value for one fish, the line the average value across fish, error bars are s.e.m.

Larvae with V3 neurons ablated did not show any gross morphological or behavioral defects, and initiated swim events at a slightly increased rate compared to control larvae (ablated: 0.93 ± 0.03 Hz, control: 0.78 ± 0.04 Hz, mean ± s.e.m., *n* = 28 ablated, 40 controls, p = 0.01, two-sample t-test). We thus sought to determine if there were subtle differences in swimming behaviors or kinematics, looking across several kinematic parameters (Fig. S5). To compare the free-swimming behavior to our results from voltage imaging and optogenetic activation, we measured the swim speed and duration for each bout, at different grating speeds. We found that control larvae continually increased their swim speed with increasing OMR grating speed (slope: 0.51 ± 0.06, mean ± s.e.m.), whereas V3 ablated larvae showed a reduced dependance of swim speed on OMR speed (slope: 0.17 ± 0.09, p = 0.0001, see Table 2 for details about the linear regression model). This finding indicates that V3 ablated larvae are less able to modulate their swim speed than are controls. Ablation of V3 neurons had no significant effect on bout duration (WT: 175 ± 3 ms, KO: 180 ± 4 ms) and very small effect on tail beat frequency (WT: 21.1 ± 0.1 Hz, KO: 21.6 ± 0.3 Hz, Fig. 6C).

## Discussion

### Voltage imaging for mapping neural dynamics in zebrafish spinal cord

Simultaneous voltage imaging from many neurons during behavior is a powerful method to characterize the repertoire of neural dynamics and to infer possible circuit mechanisms, but applications to dissecting circuit mechanisms remain sparse (Fan et al., 2020). By combining light sheet voltage imaging and fictive behavior in a virtual reality environment we identified and characterized distinct functional subpopulations in the zebrafish spinal cord. Voltage imaging was an essential tool in our studies because the underlying behaviors (tail oscillations) were far faster than Ca^2+^ imaging kinetics, and because subthreshold potentials were a critical part of the analysis.

The zebrafish spinal cord is particularly well suited to analysis by voltage imaging because its long and thin geometry coincides with the elongated camera field-of-view required for high-speed imaging; and light-sheet microscopy provides optical sectioning *in vivo* for high sensitivity and good background rejection. Further, VNR recordings and a closed-loop virtual reality environment permitted studies of naturalistic dynamics in a preparation where motion artifacts were minimal while still inducing naturalistic behavior. The VNR recording provided a means to synchronize activity measured at different focal planes, fields of view and animals.

The primary limitation in our approach is that, even with a 1 kHz recording rate, the very fast spikes of the dorsal oscillatory neurons were only detectable above noise in the most strongly expressing neurons. Further improvements in sensor speed, brightness, sensitivity, and expression are needed, along with improved high-speed imaging systems, to reliably study spiking in this population of neurons. In principle, the entire zebrafish spinal cord is accessible to light-sheet voltage imaging, and with advances in optics it may become possible to record from a substantial fraction of the neurons simultaneously (Yoon et al., 2020; Zhang et al., 2021).

### Excitatory activity in the zebrafish spinal cord

We identified three main groups of glutamatergic neurons active during slow forward swimming: oscillating neurons, dorsal non-oscillating neurons and ventral non-oscillating neurons. The oscillating neurons are most likely V0v and V2a neurons, which have been described as the major excitatory source that drive the CPG in the larval zebrafish (Ljunggren et al., 2014; McLean et al., 2007, 2008). V0v and V2a neurons have been shown to be recruited in a frequency dependent manner, and V0v neurons are inhibited at higher swimming frequencies. Since our experiments mainly induced low frequency swimming, we did not distinguish between these two classes in our analyses.

By measuring subthreshold membrane voltages, we mapped the phases of the oscillatory subpopulation across and along the spinal cord. Undulatory locomotion is generated by a linear phase gradient of motor activity and muscle contraction along the spinal cord (Grillner et al., 1993; Masino and Fetcho, 2005). Our recordings show a similar phase gradient and propagation of activity at the level of excitatory premotor interneurons.

To our knowledge, V3 neurons are the first identified population of motor-related spinal cord neurons that fire tonically instead of being phase locked to the CPG. However, the occasional phase-locked inhibition we observed in V3 neurons suggest that they can be controlled with cycle-timing precision. Such inhibition might be necessary when performing fast corrective movements or when switching to a higher frequency swim module mid-bout which would require coordinated deactivation of the slow swim circuits. V1 inhibitory neurons have been shown to provide in-phase inhibition to silence slow motor neurons at higher frequencies and could be a source for phase locked inhibition in V3 neurons. (Kimura and Higashijima, 2019; Sengupta et al., 2021). Morphological reconstruction showed that there are at least two classes of V3 neurons which can be distinguished by their descending or bifurcating axon.

Sensory input is generally received in the dorsal part of the spinal cord. Sensory gating has been described for many sensory systems, including cutaneous touch input in zebrafish (Knogler and Drapeau, 2014). A subset of the non-oscillatory dorsal neurons spiked between swim bouts and were silent during bouts (Fig. S3). We speculate that these neurons might receive sensory input that is shunted during movement. Our dataset provides a starting point to investigate the role of sensory input in spinal interneurons and their gating during behavior.

### V3 neurons as modulators of swim strength

In free-swimming fish, V3 neurons are necessary for fish to track changes in OMR grating speed. Our voltage imaging and optogenetic experiments point to a V3 function that is not along the canonical frequency-speed axis (Callahan et al., 2019; Müller and van Leeuwen, 2004; Severi et al., 2014), but rather through modulation of swim strength. This observation suggests complementary modes of CPG modulation, corresponding approximately to frequency modulation and amplitude modulation. It is not known how these different modes of modulation interact. A frequency independent effect of V3 activity on motor neurons has also recently been suggested in a study using laser ablations of V3 neurons (Wiggin et al., 2021).

Although our optogenetic experiments showed that V3 activation was sufficient to trigger swim bouts (in the presence of NMDA), our fictive swimming and ablation experiments indicated that V3 activity is not necessary for swimming. Specifically, in the fictive swimming assays, low-strength swim bouts often occurred without firing of any V3 neurons in the field of view (Fig. 3), and in free-swimming fish lacking V3 neurons, the rate of bout initiation was actually slightly higher than in sibling controls (Fig. 6). These observations suggest that V3 neurons are not a primary driver of bout initiation under physiological conditions, but that they rather serve to modulate the power once a bout is initiated and perhaps to sustain ongoing bouts.

Larval zebrafish can show motor learning over several bouts to adjust their motor output to changing environmental conditions (Ahrens et al., 2012; Kawashima et al., 2016). Such motor learning depends on serotonergic activity in the dorsal raphe nucleus, but the targets of the serotonergic modulation are not known. Our data suggests the hypothesis that serotonin may (directly or indirectly) modulate the gain of the spinal cord by acting on V3 neurons.

The tools and methods described here provide a unique window into spinal circuit dynamics during (fictive) behaviors. Measurements *in vivo* are essential for identifying naturalistic firing patterns. These approaches are expected to generalize to other spinal sensory and motor populations.

## Supporting information

Movie 1: Dynamics of oscillatory CPG neurons

Movie 2: Dynamics of non-oscillatory V3 neurons

Movie 3: Subcellular action potential propagation

## Acknowledgments

We thank S. Begum, A. Klaeger, and D. Brinks for technical assistance and J. Miller and K. Hurley for fish care. We thank E. Boyden, E. Jung, and K. Piatkevich for early access to the zArchon1 fish. U.B. and A.E.C. were supported by the Howard Hughes Medical Institute, and Office of Naval Research Vannevar Bush Faculty Fellowship and NIH grant R01MH11704201. Y.K. and S.H. were supported by National BioResource Project from the Ministry of education, Culture, Sports, Science, and Technology of Japan. F.E. was supported by National Institutes of Health (U19NS104653, R43OD024879, and 2R44OD024879), the National Science Foundation (IIS-1912293), and the Simons Foundation (SCGB 542973). T.K. and M.B.A. were supported by the Howard Hughes Medical Institute.

## Material and Methods

### Zebrafish husbandry

All zebrafish experiments were approved by the Harvard University Institutional Animal Care and Use Committee (IACUC). All larvae were kept at a 10 h dark, 14 h light cycle at 28 °C until at least 5 dpf.

### List of transgenic lines

| Name | Reference |
| --- | --- |
| *Tg(vGlut2a:Gal4)* | (Satou et al., 2013) |
| *Tg(UAS:zArchon1-GFP)* | This study |
| *Tg(vglut2a:loxP-RFP-loxP-ChR)* | This study |
| *Tg(nkx2.2b-Cre)* | This study |
| *Tg(sim1a:loxP-DsRed-loxP-DTA)* | This study |
| *Tg(hox4a/9a:Cre)* | (Kimura and Higashijima, 2019) |
| *Tg(UAS:Kaede)* | (Hatta et al., 2006) |
| *vGlut2a:GFP* | (Satou et al., 2012) |

### Zebrafish transgenesis

*Tg(vglut2a:loxP-RFP-loxP-ChR)* fish were generated using the bacterial artificial chromosome (BAC) zK145P24 as described (Kimura et al., 2006). The loxP-RFP-loxP-ChR DNA construct was generated using the loxP-RFP-loxP-GFP construct: GFP was replaced with ChR-WideReceiver (Wang et al., 2009). *Tg(nkx2*.*2b:Cre)* fish were generated using the BAC transgenic method with BAC zKP50F5. The Cre DNA construct was described in (Satou et al., 2012). *Tg(sim1a:loxP-RFP-loxP-DTA)* fish were established using the CRISPR/Cas9 knock-in method (Kimura et al., 2014) with the DNA construct described in (Kimura and Higashijima, 2019). The insertion site was set upstream of the sim1a gene with the CRISPR target sequence: GAAGCGGCCGCCGGTGAATGGG.

The *Tg(UAS:zArchon1-KGC-GFP-ER2)* line was generated using the Tol2 transposon system. The plasmid (Piatkevich et al., 2018) and tol2 mRNA were injected in 2-cell stage embryos. We crossed the injected fish with a Gal4 line to screen for the expression of green fluorescence in the brain at the F1 generation.

### Imaging setup

All voltage imaging data were acquired on a custom-built light sheet microscope (Fig. S1). Output of an 800 mW 639 nm laser (MLL-FN-639, CNI Laser) was first expanded 2x isotopically and then stretched 2x along the x-axis using an anamorphic prism pair (PS875-A, Thorlabs). The light sheet was created using a 50 mm cylindrical lens focusing along the x-axis and reimaged onto the back focal plane of a 4x long working-distance objective (Olympus XLFluor4X/340, NA 0.28) using a scan lens (50 mm, AC254-050-A-ML, Thorlabs) and tube lens (150 mm, AC508-150-A-ML, Thorlabs). Nominal specifications for the sheet are: Rayleigh length: 46 μm, sheet thickness at focus 6.1 μm, sheet width 708 μm (all values as 1/e^2^). A galvo mirror at focal distance from the scan lens allowed moving the light sheet along the z-axis. The focal plane of the imaging objective was synchronously adjusted by moving the imaging objective with a piezo actuator (PIFOC, PD72Z4CA0).

Fluorescence was acquired through a 25x objective (Olympus XPlan N 25x WMP, NA 1.05) and a 100 mm tube lens (Thorlabs TTL-100 A), resulting in an effective magnification of 13.9x. Emission light was filtered using a 664 nm long-pass filter and images were acquired on sCMOS camera (Hamamatsu, Orca Flash4) at 996.3 frames/s. Typical illumination powers were 290 mW.

Three or four z-positions were recorded per x,y-position along the tail. At the end each recording a z-stack of the zArchon1 fluorescence across the entire depth of the spinal cord was recorded to reference each recorded neuron to anatomical coordinates in the spinal cord. The zArchon1-GFP expression pattern in Fig. 1B was recorded with a custom-built two-photon excitation arm build on the same microscope.

### Sample preparation

Zebrafish embryos at 5-7 days post fertilization (dpf) were paralyzed by immersing in 1 mg/ml alpha-bungarotoxin for 1-2 minutes before mounting in 1.5% low melting point agarose on a custom-built sample holder. For ventral nerve root (VNR) recordings the agarose was removed around a few segments of the tail. The sample holder together with the electrophysiology head stage was mounted on a x-y stage holding the larva inside a stationary sample chamber containing standard fish water. The larva and electrophysiology head stage can be moved together independently of the sample chamber. This design allowed to keep the sample chamber at a fixed distance from the light-sheet objective to avoid changes in the focal plane of the light when repositioning the sample (Fig. S1).

### Electrophysiology

VNR recordings were performed as previously described (Masino and Fetcho, 2005). Recording pipettes were pulled from glass capillaries (1B150F-4, WPI) to have a ∼20 μm opening, filled with standard fish water and attached to the side of the larva by gentle suction. Data were acquired in current clamp mode using an Axon 200-B or 700-B patch clamp amplifier (Molecular Devices). The signal was low-pass filtered at 1 kHz, passed through a denoising filter to remove line noise (HumBug, Quest Scientific) and digitized at 50 kHz using custom written LabView code.

### Closed-loop virtual environment

Larvae were presented with a forward moving square grating (grating period: 2 cm, moving speed 9.9 mm/s, fish position ∼2 cm above the screen) using a video projector (Epson VS240). The light from the projector was filtered with a bandpass filter (575/40 nm) to prevent bleed-through of the projector light into the imaging channel, demagnified (f_L1_ = 200 mm) and projected onto white paper placed below the fish.

To provide closed-loop visual feedback from fictive swimming activity, we adapted the approach described in (Ahrens et al., 2012) and implemented the online feedback on a microcontroller (Teensy 3.6, PJRC). The ventral nerve root signal was sampled at 10 kHz and filtered by subtracting an exponential moving average (α = 0.005/sample). The resulting high-pass filtered signal was used to calculate the standard deviation over a moving window of 10 ms. Signals that crossed a threshold value were assigned as swim events. To calculate the swimming strength, signal values exceeding the threshold were summed over time and filtered through a high-pass filter with a time constant of 100 ms. The output of the microcontroller was sent to LabView code which applied a user-defined gain and then subtracted the scaled motion signal from the constant forward speed of the grating and generated the video output for the projector. The feedback loop ran at a rate of 100 Hz, above the video projector refresh rate (60 Hz).

To calculate the final speed signal, for each fish a constant gain scale-factor was selected so that it could follow the grating at speeds of 9.9-15.1 mm/s and at variable gain of 0.5-1.5. Four consecutive 25 s recordings of the same field of view were recorded while changing the feedback gain from 0.5 to 1.5 or the grating speed from 9.9 mm/s to 15.1 mm/s. Grating speed was kept at 0 for the first 5 s of the recording.

### Data analysis

#### VNR recordings

VNR recordings were band-pass filtered between 300 and 1000 Hz and the standard deviation over a moving 10 ms window followed by a moving average of the same window length was used to calculate the swim signal.

#### Fluorescence recordings

Using the average intensity image as a guide, masks were manually drawn around each soma in the field of view. Each frame was filtered with a median filter of 3×3 pixel and the fluorescence time-series at each pixel was high-pass filtered by subtracting a sliding median over a 201 ms window. Then PCA/ICA was performed on each cell body individually. The resulting components were inspected manually and one component per cell body was kept if it showed activity during the recording. The final fluorescence signal was calculated as *Δ*F/F = (*F*_*ICA*_(*t*)– *mean*(*F*_*ICA*_))/*mean*(*F*_*ICA*_).

To calculate the cycle-triggered average fluorescence waveform, the fluorescence was triggered on each VNR burst with a period in the range of 25 ms to 66.7 ms (14 Hz – 40 Hz). To average over cycles with different periods, individual cycles were linearly interpolated to have the same number of samples. The cycle amplitude was then calculated as the difference between the minimum and maximum value of the cycle-triggered average.

To analyze the phase relationships of oscillating neurons, only neurons with an average cycle amplitude ≥ 0.5 ΔF/F were included. To correct for apparent phase offsets between fish due to different placements of the VNR electrode, the average phase shift of all recorded cells in each FOV was set to 0.

Non-oscillating cells were defined as cells with a cycle triggered average amplitude < 0.5 ΔF/F. Only non-oscillating cells were present in the ventral spinal cord. V3 neurons were thus defined as having a cell body position in the lower 30% of the spinal cord. To quantify the difference in spike rate during vs between bouts, a bout activity index was defined as

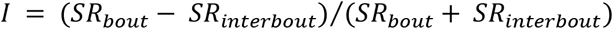

where *SR* is the spike rate. Only V3 neurons with *I* > 0 were included in the analysis.

### Sub-Nyquist action potential timing

The calculation of the action potential delay along the neurite was adapted from (Hochbaum et al., 2014). We first calculated the spike triggered average movie of each cell. Due to the rolling shutter of the sCMOS camera, this movie contained a time delay of 9.74436 μs between successive rows of pixels. We used digital interpolation to correct for this time delay. We used the spatial footprint of the neuron (see below) to trace the neurites of each neuron. We then calculated the spike waveform along the neurite by averaging the pixel values from successive 1 pixel long and 8 pixel wide regions. The resulting matrix (distance x time) was then filtered with a moving average of 40 pixels along the distance axis and a smoothing spline was fitted at each distance along the time axis (Matlab fit method ‘SmoothingSpline’, smoothing parameter 0.6). The position of the peak of the resulting function was considered the timing of the action potential maximum. For the analysis of action potential propagation, we measured the time delay along the longest neurite in the FOV from the 7 neurons that showed the strongest signal.

### Spike-based extraction of neuronal morphology

To visualize the morphology of individual V3 neurons from the voltage imaging data, we generated spike triggered average movies and the corresponding spike triggered average fluorescence trace. We then calculated the correlation coefficient of the timeseries of each pixel and the average spike waveform. The resulting image was filtered with a 2D median filter with a radius of 2 pixels to remove noise. This approach gave an image of the single cell body and proximal neurites.

### Kaede conversion and single-cell morphology

The Kaede photoconversion protocol was adopted from (Callahan et al., 2019). Zebrafish embryos at 4 dpf with genotype *Tg(vGlut2a:Gal4; UAS:Kaede)* were anesthetized with 0.02% Tricaine and mounted in 1.5% low melting-point agarose. One or two Kaede-expressing neurons at the ventral edge of the spinal cord were targeted for conversion. Photoconversion was performed with a 405 nm laser at maximum intensity using the bleach function on a Zeiss LSM-710 confocal microscope equipped with a 40x W Plan-Apochromat objective. Pulses of scanning over the cell body were repeated in 1 s intervals interleaved with imaging the converted and unconverted fluorescence channels until the signal of the photoconverted protein reached a steady state. Larvae were left in fish water overnight to allow photoconverted protein to diffuse along the axon before remounting for imaging the next day. Neuron morphology was segmented using the simple neurite tracer plugin in ImageJ (Longair et al., 2011).

### Channelrhodopsin stimulation

Zebrafish embryos at 5-6 dpf with genotype *Tg(vglut2a:loxP-DsRed-loxP-ChR; nkx2*.*2b:Cre-mCherry-NLS)* expressing ChR WideReceiver in the V3 neurons were spinalized at the level of the swim bladder and mounted in 1.5% low melting-point agarose. Agarose was removed around the tail and NMDA was added to a final concentration of 50 μM. After 1 min. incubation time, the tail motion was recorded at a frame rate of 200 Hz and the angle was calculated in real time using custom LabView code. The standard deviation over a 10 ms window of the tail angle was used as proxy for the larva’s swimming speed. The entire tail was illuminated with 470 nm blue light at 1.3-5.9 mW/mm^2^ with a 3 s on 3 s off cycle for 1 min. This intensity was sufficient to saturate the response of the ChR WideReceiver (Umeda et al., 2013). The NMDA concentration was then increased to 75 μM and 100 μM respectively and the experiment repeated. Fish were excluded if none of the applied NMDA concentrations resulted in clear tail oscillations or if there were not at least 5 swim bouts in the light-off condition (9 excluded out of 19 fish).

### Free swimming experiments

To create fish lacking V3 neurons, either *Tg(sim1a:*|*DsRed*|*-DTA)*^*+/-*^ fish were crossed to *Tg(hox4a/9a:Cre)*^*+/-*^ fish or *Tg(sim1a:*|*DsRed*|*-DTA*^*+/-*^; *hox4a/9aCre*^*+/-*^) fish were out-crossed to wildtype adults. The resulting offspring contained a mix of larvae containing the sim1a:|DsRed|-DTA transgene with or without the Cre recombinase. Larvae were pre-screened for the presence of the *Tg(sim1a:*|*DsRed*|*-DTA)* transgene by strong DsRed expression in the brain under low magnification. Experiments were performed blind to Cre expression and hence whether the fish contained V3 neurons. After the experiment, spinal cord expression of DsRed was verified under high magnification to assign ‘KO’ (larvae lacking V3 neurons) or ‘WT’ (larvae expressing DsRed in V3 neurons) genotypes.

To verify the ablations of V3 neurons, we crossed *Tg(sim1a:*|*DsRed*|*-DTA*^*+/-*^; *hox4a/9aCre*^*+/-*^) to *Tg(vGlut2a:GFP)* (Satou et al., 2012) and acquired confocal images (Olympus FV1000) to verify the absence of V3 neurons in the ventral spinal cord (Fig. 6A)

Larvae at 5-6 dpf were placed in a 76.2 × 15.7 mm arena cast from 3% agarose. The arena was placed on top of diffusive paper (Rosco Cinegel 3000) and illuminated from below with an array of IR LEDs. A video projector (Apeman 5000L) was used to project a grating (spatial period 7.7 mm, rgb values: 0 0 0 or 0 1 1) onto the diffusive paper via a cold mirror. Larva motion was recorded at 300 Hz with an IR-sensitive monochrome camera (Flir, GS3-U3-41C6NIR-C) equipped with a long-pass filter and a photo objective (Sigma Zoom, 18-200 mm). Each larva received 10 minutes of stationary grating followed by 5 s moving grating and 5 s stationary grating for 10 minutes. The order of grating speeds presented was shuffled for each trial and if a larva reached the end of the arena before the end of the 5 s period, the grating was stopped, and the sequence proceeded into the 5s static period. All stimulus presentation and live tracking of the fish were done with the Stytra software package (Štih et al., 2019).

To exclude false positive swim events due to failed video tracking, we excluded all identified bouts with a standard deviation of the instantaneous cycle frequency > 0.07. Fish where the fraction of excluded bouts exceeded 10% were excluded from the analysis (9 excluded out of 77 fish).

### Statistical significance testing

#### ChR activation experiments

We sought to test whether blue light optogenetic stimulation (LED) affected bout duration, swim strength or tail beat frequency. Each fish was tested multiple times with and without illumination, and we sought to control for inter-animal differences. We fitted a linear mixed model using the matlab function `fitlme`, and p-values were obtained using `anova` on the resulting model. After correcting for multiple hypothesis testing, a significant effect of LED illumination was set at p ≤ 0.0167.

|  | *Estimate ($\log_{10}(\text{ms})$)* | *SE* | *FStat* | *pValue* |
| --- | --- | --- | --- | --- |
| (Intercept) | 2.7116 | 0.12176 | 495.97 | 1.1074e-84 |
| LED | 0.22015 | 0.08658 | 6.4657 | 0.011198 |

$\text{SwimStrength} \sim 1 + \text{LED} + (1 + \text{LED} \mid \text{fishID})$
|  | *Estimate (a.u.)* | *SE* | *FStat* | *pValue* |
| --- | --- | --- | --- | --- |
| (Intercept) | 0.05041 | 0.0087851 | 32.926 | 1.445e-08 |
| LED | 0.012388 | 0.0040424 | 9.3918 | 0.0022663 |

$\text{Frequency} \sim 1 + \text{LED} + (1 + \text{LED} \mid \text{fishID})$
|  | *Estimate (Hz)* | *SE* | *FStat* | *pValue* |
| --- | --- | --- | --- | --- |
| (Intercept) | 12.836 | 0.64597 | 394.85 | 1.4986e-69 |
| LED | 0.55402 | 0.26971 | 4.2196 | 0.040344 |

#### Free swimming experiments

In the free swimming experiments, we sought to test whether the genotype had a significant effect on swim speed, bout duration or tail beat frequency as a response to swimming at different OMR grating speeds. We fitted a linear mixed model using the matlab function `fitlme`, and p-values were obtained using `anova` on the resulting model. After correcting for multiple hypothesis testing, significance for either genotype or the genotype:omr interaction was set at p ≤ 0.0167.

|  | *Estimate (mm/s)* | *SE* | *FStat* | *pValue* |
| --- | --- | --- | --- | --- |
| (Intercept) | 6.8846 | 0.25526 | 727.47 | 2.7028e-72 |
| omr | 0.51001 | 0.052955 | 92.757 | 1.3644e-18 |
| genotype | 0.901 | 0.43914 | 4.2096 | 0.041358 |
| omr:genotype | -0.34485 | 0.087244 | 15.624 | 0.00010362 |

duration ~ 1 + omr\*genotype + (1 + omr | fishId)
|  | *Estimate (s)* | *SE* | *FStat* | *pValue* |
| --- | --- | --- | --- | --- |
| (Intercept) | 0.1555 | 0.0033605 | 2141.2 | 1.2667e-116 |
| omr | 0.0044722 | 0.00042461 | 110.93 | 2.4818e-21 |
| genotype | 0.004082 | 0.0056441 | 0.52306 | 0.47029 |
| omr:genotype | -0.00079233 | 0.00069955 | 1.2828 | 0.25858 |

frequency ~ 1 + omr\*genotype + (1 + omr | fishId)
|  | *Estimate (Hz)* | *SE* | *FStat* | *pValue* |
| --- | --- | --- | --- | --- |
| (Intercept) | 19.996 | 0.17552 | 12979 | 2.7737e-200 |
| omr | 0.24289 | 0.015327 | 251.14 | 1.9345e-38 |
| genotype | 0.74014 | 0.29456 | 6.3135 | 0.012687 |
| omr:genotype | -0.083002 | 0.025251 | 10.805 | 0.0011751 |

---

**Supplementary figure 1.**
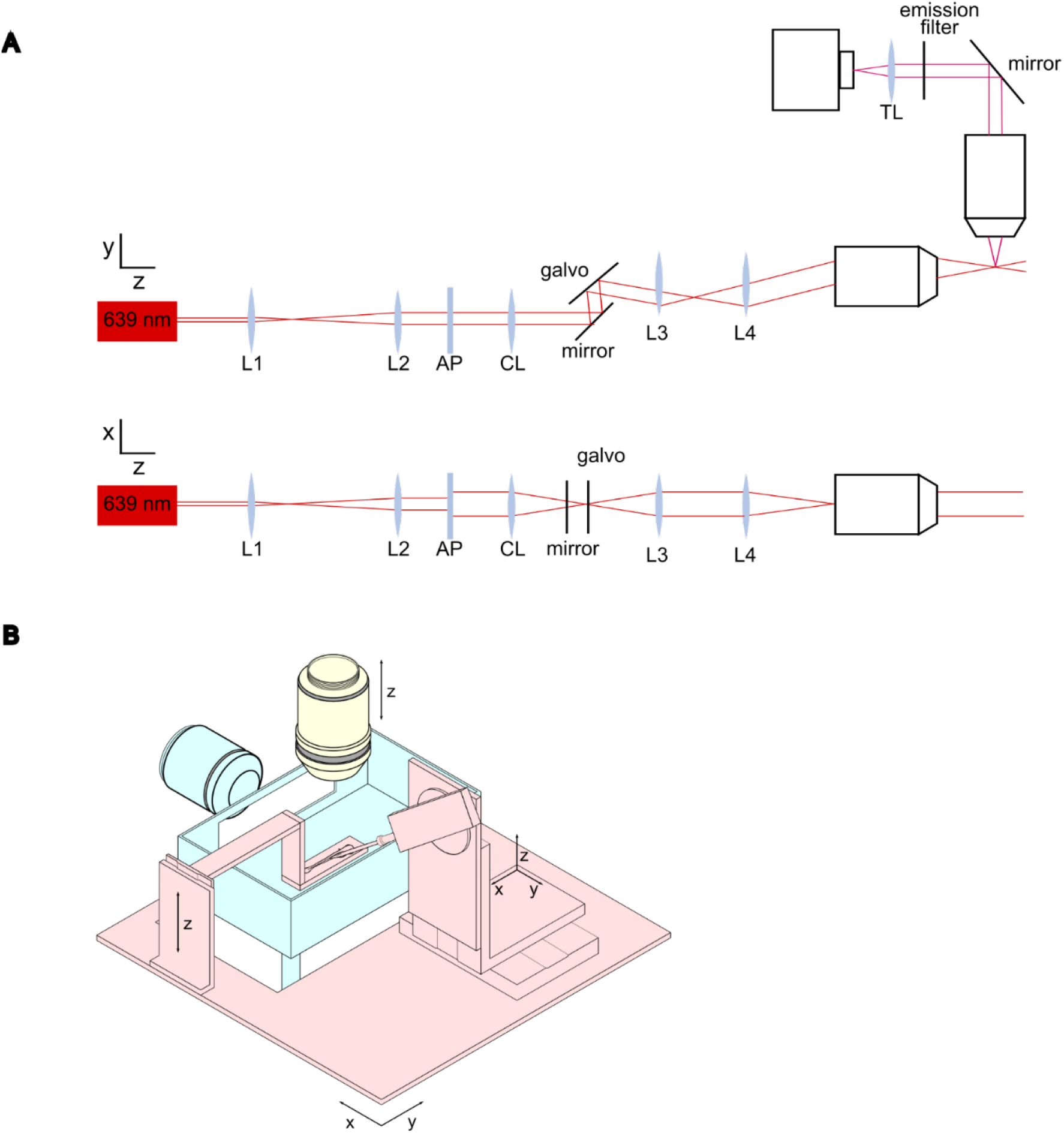
Light-sheet voltage imaging setup. (A) Schematic of the optical setup. Side and top view of the excitation beam path showing the beam expander (L1 and L2), anamorphic prism pair to expand the beam along the x-axis, cylindrical lens (Cl), folding mirror and galvo mirror to position the light sheet along the y-axis, scan lens (L3), tube lens (L4) and illumination objective. The emission path contains the imaging objective, folding mirror, emission filter, tube lens (TL) and camera. See Materials and Methods for specifications about the individual components. (B) Sample holder and imaging chamber. Similarly colored elements can move together. Red: sample holder and electrophysiology headstage on x-y stage. Yellow: imaging objective. Blue: sample chamber and illumination objective (fixed).

**Supplementary figure 2.**
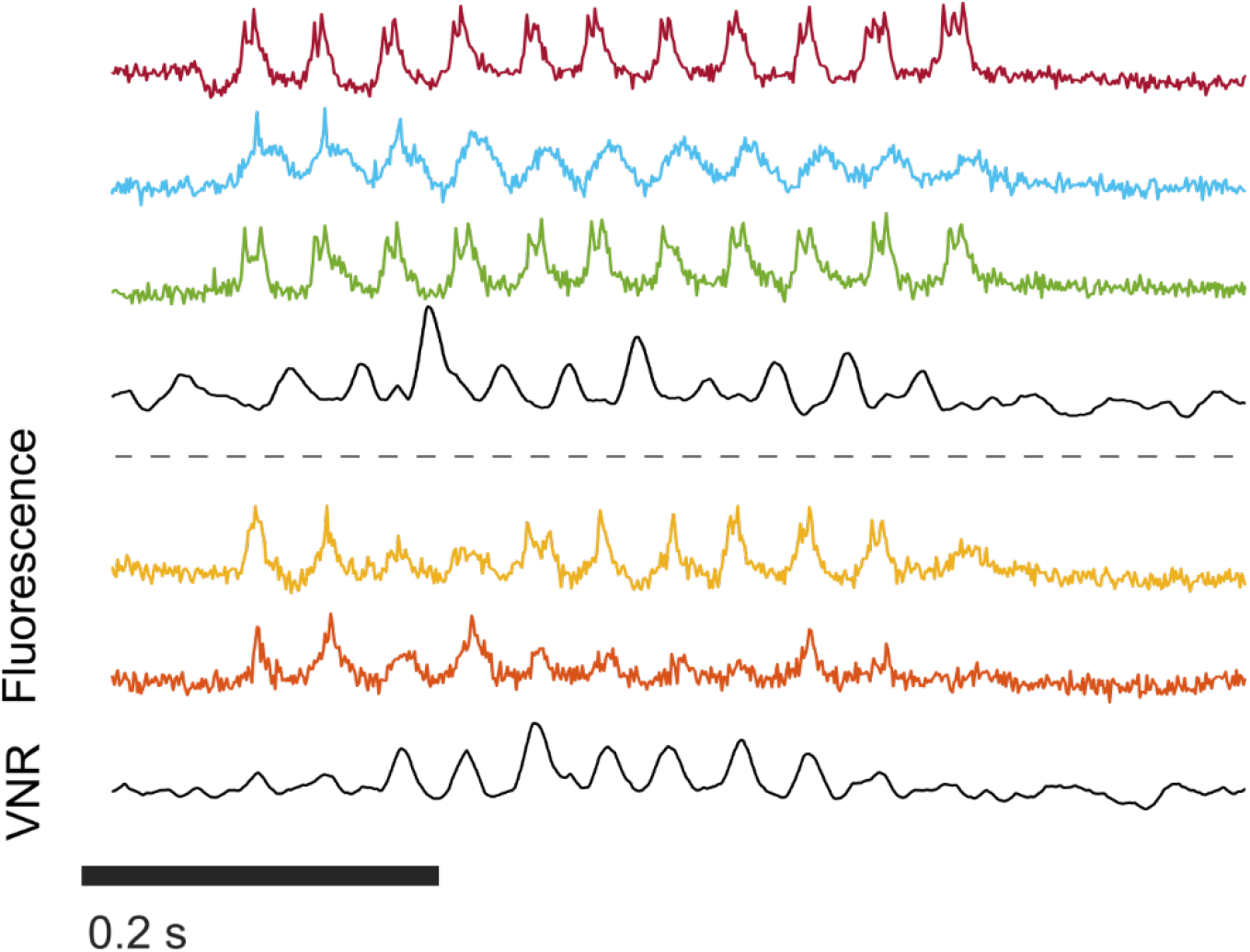
High SNR recordings show spiking in oscillating neurons. High SNR optical voltage recordings and corresponding VNR signals from oscillating neurons during a swim bout. Small spikes were visible on top of the larger subthreshold depolarizations. Dashed line separates recordings from different animals.

**Supplementary figure 3.**
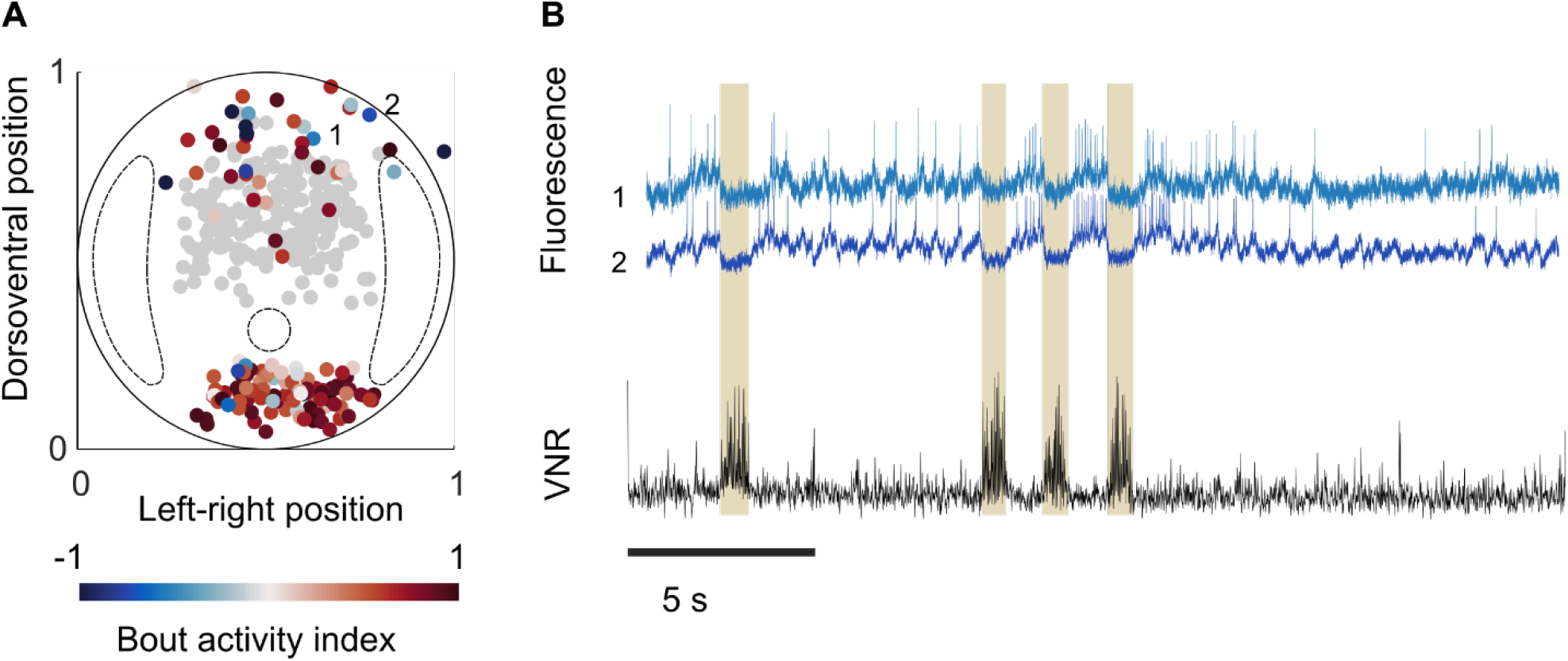
Distribution of non-oscillating neurons active during bouts and interbout intervals. (A) Transverse view showing cell body positions of non-oscillating neurons color coded by bout activity index. Oscillating neurons are in gray. Fluorescence traces of numbered neurons are plotted in (B). (B) Fluorescence traces of two dorsal neurons imaged simultaneously showing distinct spiking during interbout intervals and inhibition during swim bouts.

**Supplementary figure 4.**
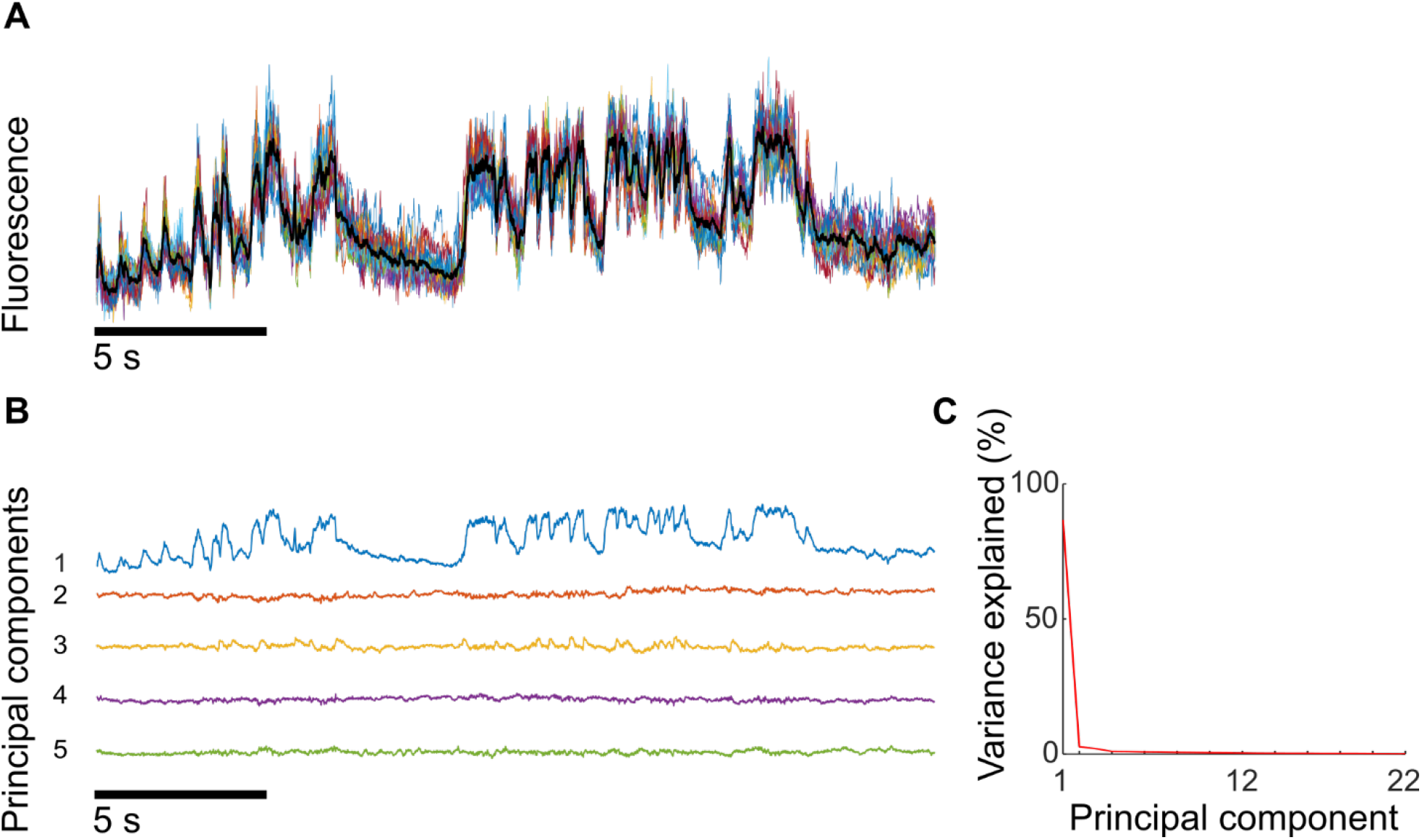
Subthreshold voltages of V3 neurons vary primarily along a single dimension. (A) Individual and average subthreshold voltage (spikes removed) of 22 simultaneously recorded V3 neurons. Subthreshold fluctuations were highly synchronized between all neurons. (B) The first five principal components show that most of the subthreshold variance is captured in the first principal component. (C) Percent of variance explained by each of the 22 principal components of the subthreshold voltages.

**Supplementary figure 5.**
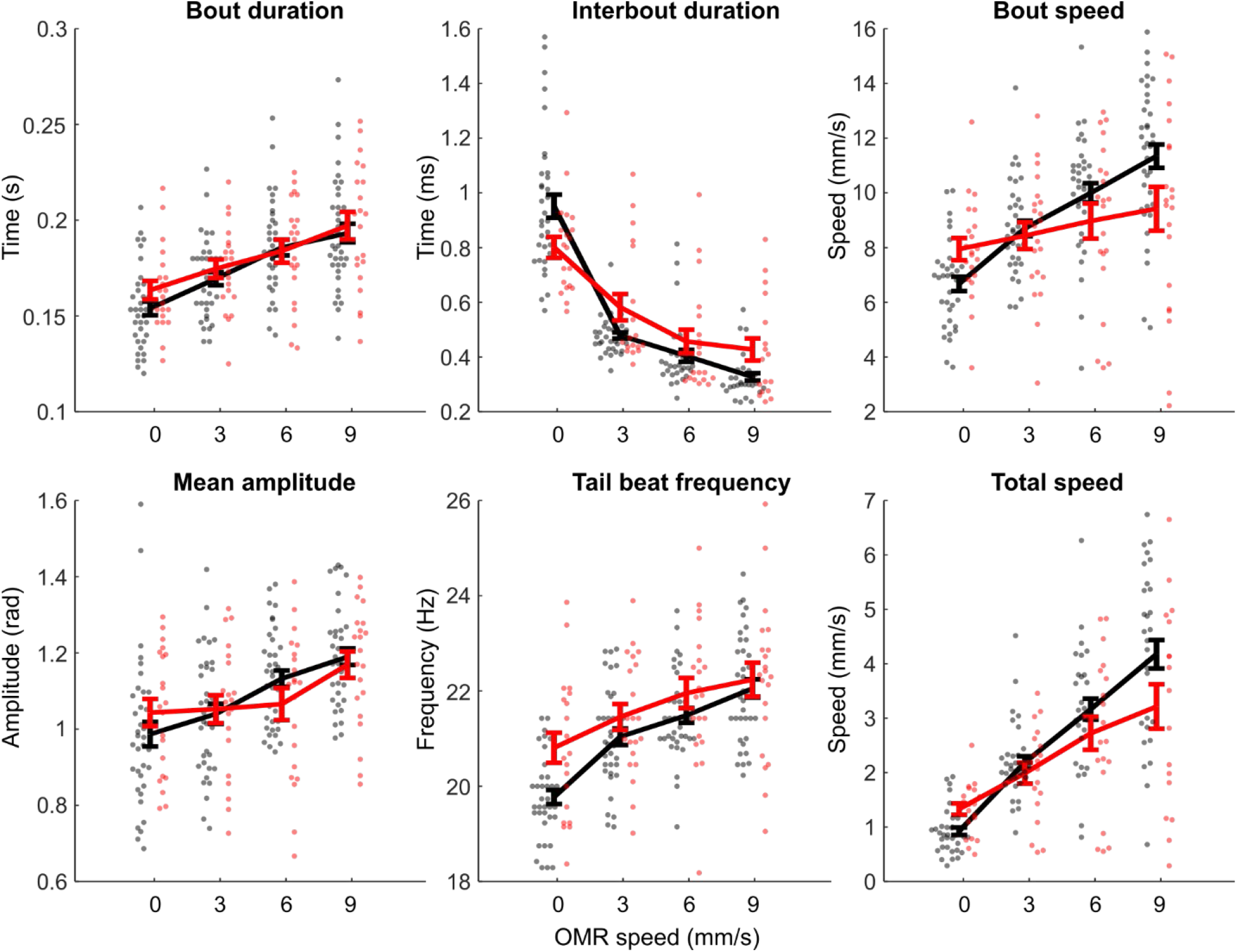
Kinematics of free-swimming V3 ablated and control larvae. Red: V3 ablated. Black: clutch-matched controls. All parameters were measured as a function of OMR grating speed. Each dot represents the median value for one fish, the line the average value across fish, error bars are s.e.m.

## Notes

### Competing Interest Statement

The authors have declared no competing interest.

## References

Abdelfattah, A.S., Kawashima, T., Singh, A., Novak, O., Liu, H., Shuai, Y., Huang, Y.-C., Campagnola, L., Seeman, S.C., Yu, J., et al. (2019). Bright and photostable chemigenetic indicators for extended in vivo voltage imaging. Science 365, 699.

Adam, Y., Kim, J.J., Lou, S., Zhao, Y., Xie, M.E., Brinks, D., Wu, H., Mostajo-Radji, M.A., Kheifets, S., and Parot, V. (2019). Voltage imaging and optogenetics reveal behaviour-dependent changes in hippocampal dynamics. Nature 569, 413.

Ahrens, M.B., Li, J.M., Orger, M.B., Robson, D.N., Schier, A.F., Engert, F., and Portugues, R. (2012). Brain-wide neuronal dynamics during motor adaptation in zebrafish. Nature 485, 471–477.

Ahrens, M.B., Huang, K.-H., Narayan, S., Mensh, B.D., and Engert, F. (2013). Two-photon calcium imaging during fictive navigation in virtual environments. Front. Neural Circuits 7.

Ampatzis, K., Song, J., Ausborn, J., and El Manira, A. (2014). Separate microcircuit modules of distinct v2a interneurons and motoneurons control the speed of locomotion. Neuron 83, 934–943.

Borowska, J., Jones, C.T., Zhang, H., Blacklaws, J., Goulding, M., and Zhang, Y. (2013). Functional Subpopulations of V3 Interneurons in the Mature Mouse Spinal Cord. J Neurosci 33, 18553–18565.

Bruno, A.M., Frost, W.N., and Humphries, M.D. (2015). Modular Deconstruction Reveals the Dynamical and Physical Building Blocks of a Locomotion Motor Program. Neuron 86, 304–318.

Budick, S.A., and O’Malley, D.M. (2000). Locomotor repertoire of the larval zebrafish: swimming, turning and prey capture. J. Exp. Biol 203, 2565–2579.

Cai, C., Friedrich, J., Singh, A., Eybposh, M.H., Pnevmatikakis, E.A., Podgorski, K., and Giovannucci, A. (2021). VolPy: Automated and scalable analysis pipelines for voltage imaging datasets. PLOS Computational Biology 17, e1008806.

Callahan, R.A., Roberts, R., Sengupta, M., Kimura, Y., Higashijima, S., and Bagnall, M.W. (2019). Spinal V2b neurons reveal a role for ipsilateral inhibition in speed control. ELife 8, e47837.

Chien, M.-P., Brinks, D., Testa-Silva, Guilherme, Tian, He, Brooks F. Phil, Adam, Y., Bloxham, W., Gmeiner, Benjamin, Kheifets, S., and Cohen, A.E. (2021). Photoactivated voltage imaging in tissue with an Archaerhodopsin-derived reporter. Science Advances 7, eabe3216.

Chopek, J.W., Nascimento, F., Beato, M., Brownstone, R.M., and Zhang, Y. (2018). Sub-populations of Spinal V3 Interneurons Form Focal Modules of Layered Pre-motor Microcircuits. Cell Rep 25, 146–156.e3.

Danner, S.M., Zhang, H., Shevtsova, N.A., Borowska-Fielding, J., Deska-Gauthier, D., Rybak, I.A., and Zhang, Y. (2019). Spinal V3 Interneurons and Left–Right Coordination in Mammalian Locomotion. Front. Cell. Neurosci. 13.

Fan, L.Z., Kheifets, S., Böhm, U.L., Wu, H., Piatkevich, K.D., Xie, M.E., Parot, V., Ha, Y., Evans, K.E., Boyden, E.S., et al. (2020). All-Optical Electrophysiology Reveals the Role of Lateral Inhibition in Sensory Processing in Cortical Layer 1. Cell 180, 521–535.e18.

Gong, Y., Huang, C., Li, J.Z., Grewe, B.F., Zhang, Y., Eismann, S., and Schnitzer, M.J. (2015). High-speed recording of neural spikes in awake mice and flies with a fluorescent voltage sensor. Science 350, 1361– 1366.

Goulding, M. (2009). Circuits controlling vertebrate locomotion: moving in a new direction. Nat. Rev. Neurosci 10, 507–518.

Grillner, S., and Manira, A.E. (2015). The intrinsic operation of the networks that make us locomote. Current Opinion in Neurobiology 31, 244–249.

Grillner, S., Matsushima, T., Wadden, T., Tegnér, J., El Manira, A., and Wallén, P. (1993). The neurophysiological bases of undulatory locomotion in vertebrates. Seminars in Neuroscience 5, 17–27.

Hale, M.E., Ritter, D.A., and Fetcho, J.R. (2001). A confocal study of spinal interneurons in living larval zebrafish. J. Comp. Neurol 437, 1–16.

Hatta, K., Tsujii, H., and Omura, T. (2006). Cell tracking using a photoconvertible fluorescent protein. Nat Protoc 1, 960–967.

Higashijima, S.-I., Schaefer, M., and Fetcho, J.R. (2004). Neurotransmitter properties of spinal interneurons in embryonic and larval zebrafish. Journal of Comparative Neurology 480, 19–37.

Hochbaum, D.R., Zhao, Y., Farhi, S., Klapoetke, N., Werley, C.A., Kapoor, V., Zou, P., Kralj, J.M., Maclaurin, D., Smedemark-Margulies, N., et al. (2014). All-optical electrophysiology in mammalian neurons using engineered microbial rhodopsins. Nat.Methods 11, 825–833.

Kawashima, T., Zwart, M.F., Yang, C.-T., Mensh, B.D., and Ahrens, M.B. (2016). The Serotonergic System Tracks the Outcomes of Actions to Mediate Short-Term Motor Learning. Cell 167, 933–946.e20.

Kimura, Y., and Higashijima, S.-I. (2019). Regulation of locomotor speed and selection of active sets of neurons by V1 neurons. Nat Commun 10, 2268.

Kimura, Y., Okamura, Y., and Higashijima, S. (2006). alx, a Zebrafish Homolog of Chx10, Marks Ipsilateral Descending Excitatory Interneurons That Participate in the Regulation of Spinal Locomotor Circuits. J. Neurosci. 26, 5684–5697.

Kimura, Y., Hisano, Y., Kawahara, A., and Higashijima, S. (2014). Efficient generation of knock-in transgenic zebrafish carrying reporter/driver genes by CRISPR/Cas9-mediated genome engineering. Scientific Reports 4, srep06545.

Knogler, L.D., and Drapeau, P. (2014). Sensory gating of an embryonic zebrafish interneuron during spontaneous motor behaviors. Front Neural Circuits 8, 121.

Ljunggren, E.E., Haupt, S., Ausborn, J., Ampatzis, K., and El Manira, A. (2014). Optogenetic activation of excitatory premotor interneurons is sufficient to generate coordinated locomotor activity in larval zebrafish. J. Neurosci. 34, 134–139.

Longair, M.H., Baker, D.A., and Armstrong, J.D. (2011). Simple Neurite Tracer: open source software for reconstruction, visualization and analysis of neuronal processes. Bioinformatics 27, 2453–2454.

Marques, J.C., Lackner, S., Félix, R., and Orger, M.B. (2018). Structure of the Zebrafish Locomotor Repertoire Revealed with Unsupervised Behavioral Clustering. Curr. Biol.

Masino, M.A., and Fetcho, J.R. (2005). Fictive swimming motor patterns in wild type and mutant larval zebrafish. J. Neurophysiol. 93, 3177–3188.

McDearmid, J.R., and Drapeau, P. (2006). Rhythmic Motor Activity Evoked by NMDA in the Spinal Zebrafish Larva. Journal of Neurophysiology 95, 401–417.

McLean, D.L., Fan, J., Higashijima, S., Hale, M.E., and Fetcho, J.R. (2007). A topographic map of recruitment in spinal cord. Nature 446, 71–75.

McLean, D.L., Masino, M.A., Koh, I.Y.Y., Lindquist, W.B., and Fetcho, J.R. (2008). Continuous shifts in the active set of spinal interneurons during changes in locomotor speed. Nat Neurosci 11, 1419–1429.

Menelaou, E., and McLean, D.L. (2012). A Gradient in Endogenous Rhythmicity and Oscillatory Drive Matches Recruitment Order in an Axial Motor Pool. J. Neurosci. 32, 10925–10939.

Müller, U.K., and van Leeuwen, J.L. (2004). Swimming of larval zebrafish: ontogeny of body waves and implications for locomotory development. Journal of Experimental Biology 207, 853–868.

Piatkevich, K.D., Jung, E.E., Straub, C., Linghu, C., Park, D., Suk, H.-J., Hochbaum, D.R., Goodwin, D., Pnevmatikakis, E., Pak, N., et al. (2018). A robotic multidimensional directed evolution approach applied to fluorescent voltage reporters. Nature Chemical Biology 14, 352–360.

Piatkevich, K.D., Bensussen, S., Tseng, H., Shroff, S.N., Lopez-Huerta, V.G., Park, D., Jung, E.E., Shemesh, O.A., Straub, C., and Gritton, H.J. (2019). Population imaging of neural activity in awake behaving mice. Nature 1–5.

Portugues, R., and Engert, F. (2011). Adaptive locomotor behavior in larval zebrafish. Front. Syst. Neurosci. 5, 72.

Satou, C., Kimura, Y., and Higashijima, S. (2012). Generation of multiple classes of V0 neurons in zebrafish spinal cord: progenitor heterogeneity and temporal control of neuronal diversity. J Neurosci 32, 1771–1783.

Satou, C., Kimura, Y., Hirata, H., Suster, M.L., Kawakami, K., and Higashijima, S. (2013). Transgenic tools to characterize neuronal properties of discrete populations of zebrafish neurons. Development 140, 3927–3931.

Sengupta, M., Daliparthi, V., Roussel, Y., Bui, T.V., and Bagnall, M.W. (2021). Spinal V1 neurons inhibit motor targets locally and sensory targets distally. Current Biology.

Severi, K.E., Portugues, R., Marques, J.C., O’Malley, D.M., Orger, M.B., and Engert, F. (2014). Neural Control and Modulation of Swimming Speed in the Larval Zebrafish. Neuron 83, 692–707.

Štih, V., Petrucco, L., Kist, A.M., and Portugues, R. (2019). Stytra: An open-source, integrated system for stimulation, tracking and closed-loop behavioral experiments. PLOS Computational Biology 15, e1006699.

Tomina, Y., and Wagenaar, D.A. (2017). A double-sided microscope to realize whole-ganglion imaging of membrane potential in the medicinal leech. ELife 6, e29839.

Umeda, K., Shoji, W., Sakai, S., Muto, A., Kawakami, K., Ishizuka, T., and Yawo, H. (2013). Targeted expression of a chimeric channelrhodopsin in zebrafish under regulation of Gal4-UAS system. Neuroscience Research 75, 69–75.

Villette, V., Chavarha, M., Dimov, I.K., Bradley, J., Pradhan, L., Mathieu, B., Evans, S.W., Chamberland, S., Shi, D., Yang, R., et al. (2019). Ultrafast Two-Photon Imaging of a High-Gain Voltage Indicator in Awake Behaving Mice. Cell 179, 1590–1608.e23.

Wahlstrom-Helgren, S., Montgomery, J.E., Vanpelt, K.T., Biltz, S.L., Peck, J.H., and Masino, M.A. (2019). Glutamate receptor subtypes differentially contribute to optogenetically-activated swimming in spinally-transected zebrafish larvae. Journal of Neurophysiology.

Wang, H., Sugiyama, Y., Hikima, T., Sugano, E., Tomita, H., Takahashi, T., Ishizuka, T., and Yawo, H. (2009). Molecular determinants differentiating photocurrent properties of two channelrhodopsins from chlamydomonas. J Biol Chem 284, 5685–5696.

Wiggin, T.D., Montgomery, J.E., Brunick, A.J., Peck, J.H., and Masino, M.A. (2021). V3 Interneurons Regulate Locomotor Vigor by Recruitment of Spinal Motor Neurons During Fictive Swimming in Larval Zebrafish. BioRxiv 2021.03.03.433646.

Xie, M.E., Adam, Y., Fan, L.Z., Böhm, U.L., Kinsella, I., Zhou, D., Rozsa, M., Singh, A., Svoboda, K., Paninski, L., et al. (2021). High-fidelity estimates of spikes and subthreshold waveforms from 1-photon voltage imaging in vivo. Cell Reports 35, 108954.

Yoon, Y.-G., Wang, Z., Pak, N., Park, D., Dai, P., Kang, J.S., Suk, H.-J., Symvoulidis, P., Guner-Ataman, B., Wang, K., et al. (2020). Sparse decomposition light-field microscopy for high speed imaging of neuronal activity. Optica 7.

Zhang, Y., Narayan, S., Geiman, E., Lanuza, G.M., Velasquez, T., Shanks, B., Akay, T., Dyck, J., Pearson, K., Gosgnach, S., et al. (2008). V3 spinal neurons establish a robust and balanced locomotor rhythm during walking. Neuron 60, 84–96.

Zhang, Z., Bai, L., Cong, L., Yu, P., Zhang, T., Shi, W., Li, F., Du, J., and Wang, K. (2021). Imaging volumetric dynamics at high speed in mouse and zebrafish brain with confocal light field microscopy. Nat Biotechnol 39, 74–83.

